# Revealing the atomic and electronic mechanism of human manganese superoxide dismutase product inhibition

**DOI:** 10.1101/2024.01.26.577433

**Authors:** Jahaun Azadmanesh, Katelyn Slobodnik, Lucas R. Struble, William E. Lutz, Leighton Coates, Kevin L. Weiss, Dean A. A. Myles, Thomas Kroll, Gloria E. O. Borgstahl

## Abstract

Human manganese superoxide dismutase (MnSOD) is a crucial oxidoreductase that maintains the vitality of mitochondria by converting O_2_^●-^ to O_2_ and H_2_O_2_ with proton-coupled electron transfers (PCETs). Since changes in mitochondrial H_2_O_2_ concentrations are capable of stimulating apoptotic signaling pathways, human MnSOD has evolutionarily gained the ability to be highly inhibited by its own product, H_2_O_2_. A separate set of PCETs is thought to regulate product inhibition, though mechanisms of PCETs are typically unknown due to difficulties in detecting the protonation states of specific residues that coincide with the electronic state of the redox center. To shed light on the underlying mechanism, we combined neutron diffraction and X-ray absorption spectroscopy of the product-bound, trivalent, and divalent states to reveal the all-atom structures and electronic configuration of the metal. The data identifies the product-inhibited complex for the first time and a PCET mechanism of inhibition is constructed.

## INTRODUCTION

About 25% of known enzymes are oxidoreductases that catalyze the electron transfers that life depends on^1^. Essential biological processes such as energy generation and DNA synthesis rely on these enzymes, and dysfunction results in a wide range of diseases^2–6^. Oxidoreductases couple the transfer of electrons to the transfer of protons in a process called proton-coupled electron transfer (PCET) to accelerate redox reactions to the rates needed for life^1,7,8^. The biochemistry behind enzymes utilizing PCETs is poorly understood as it requires a precise definition of the proton donors/acceptors coordinated with the electron transfer steps^7,9,10^. The definition of these fundamental biochemical reactions is important not only for understanding disease states but also for designing therapeutic interventions relying on PCETs such as irradiation protectants and industrial applications such as electrochemical biosensors^11–14^.

Of particular note are oxidoreductases that regulate the concentration of reactive oxygen species (ROS) in cells through PCET-mediated redox reactions. ROS levels mediate mitophagy and programmed cell death. Dysfunction of oxidoreductases responsible for limiting ROS concentrations contributes to cardiovascular disease, neurological disease, and cancer progression^15,16^. Human manganese superoxide dismutase (MnSOD) is an oxidoreductase found in the mitochondrial matrix that decreases O_2_^●--^ concentrations using PCET reactions. MnSOD eliminates O_2_^●-^ by oxidation to O_2_ with a trivalent Mn ion (*k*_1_) and reduction to H_2_O_2_ with a divalent Mn ion (*k*_2_) in two half reactions that restore the trivalent Mn ion^17,18^. MnSOD is the only means that the mitochondrial matrix has to lower O_2_^●-^ levels to avoid macromolecular damage and is essential for preserving mitochondrial function^19^.

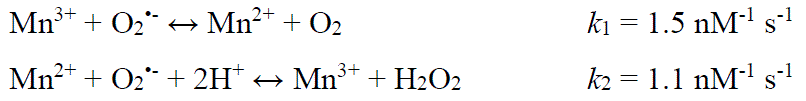

Endogenous O_2_^●-^ is produced from electrons leaking from the mitochondrial electron transport chain and necessitates MnSODs ability to eliminate O_2_^●-20,21^. Abnormalities in MnSOD activity, therefore, compromise mitochondrial function and lead to disease. In particular, genetic aberrations of MnSOD are associated with several cancer types, as noted in curated databases such as UniProt and ClinGen, with breast and prostate cancers being the most frequent^22,23^. Polymorphisms of MnSOD are a predictor for deficient cardiovascular function^24^. These disease states suggest the high reaction rate and efficiency (*k*_cat_/K_m_ > ∼ 10^9^ M^-1^ s^-1^) of MnSOD’s PCETs are key to preserving health^18,25^.

MnSOD relays protons to the active site for PCET catalysis through its hydrogen bond network (dashed blue lines, **Fig. 1a**) that extends to a neighboring subunit and is precisely packed and oriented with the aid of neighboring hydrophobic residues^25–27^. The Mn ion is bound by His26, His74, His163, Asp159, and a solvent molecule denoted as WAT1. The hydrogen bond network extends from WAT1 through Asp159, Gln143, Tyr34, another solvent molecule denoted as WAT2, His30, and Tyr166 from the neighboring subunit. Glu162 contributes to the net charge of the active site and stabilizes the dimeric interface^28,29^. Hydrophobic residues Trp123, Trp161, and Phe66 hold the hydrogen bonding atoms of Asp159, WAT1, Gln143, and Tyr34 close together^30^. Our previous work detailed changes in protonations and hydrogen bonding between Mn^3+^SOD and Mn^2+^SOD, giving insight into how the active site metal redox state alters the pK_a_s of nearby amino acids^31^.

**Fig. 1:**
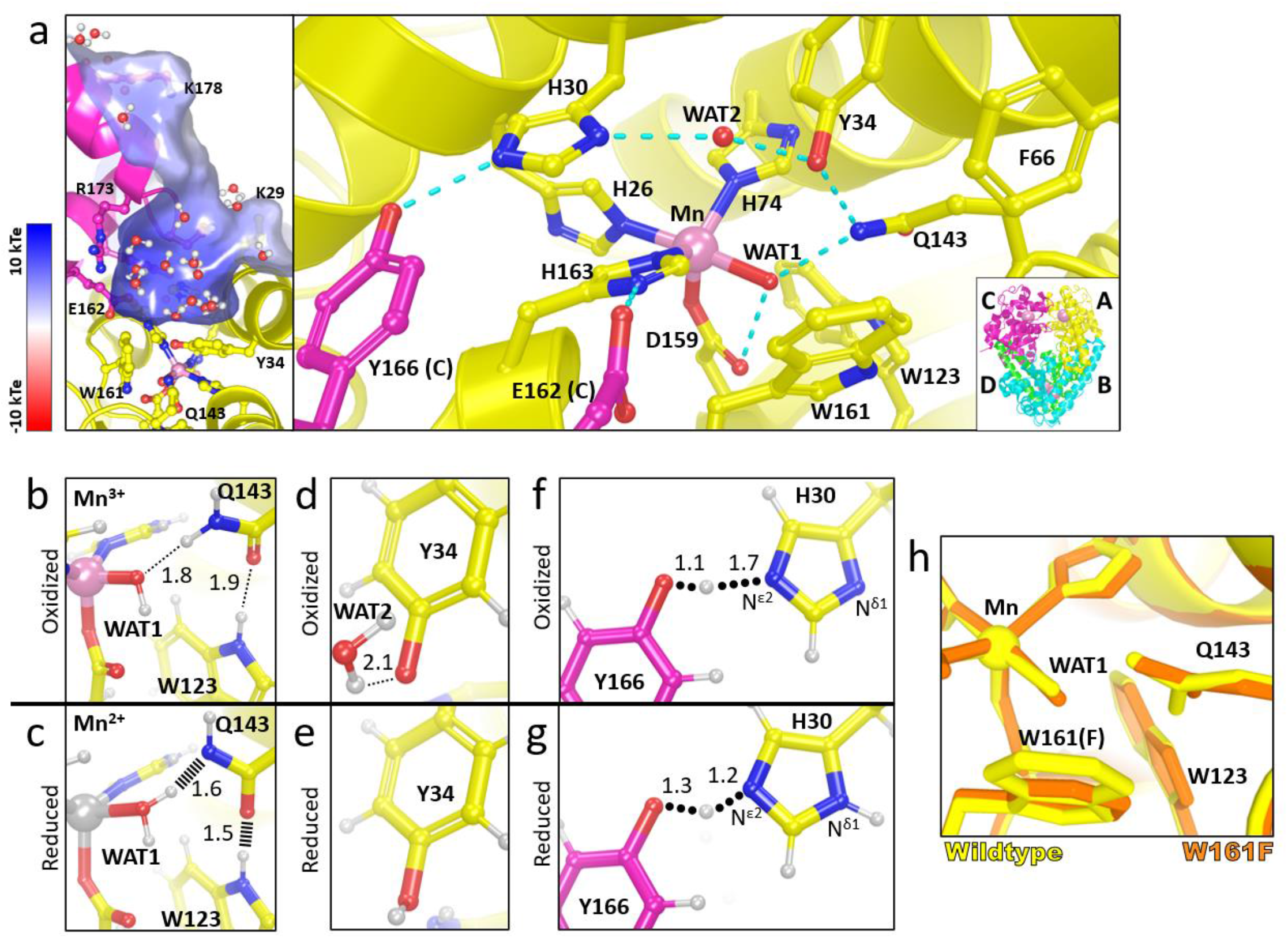
Structure of human wildtype MnSOD and protonation changes coupled to oxidation states. a. The active sites of tetrameric MnSOD are in a positively charged cavity between two subunits. Blue dashes indicate hydrogen bonds. The inset indicates chain identity where the dimeric crystallographic asymmetric unit is composed of chains A and B while C and D are generated by symmetry. **b, c** Room temperature neutron structures of Mn^3+^SOD and Mn^2+^SOD show changes in protonation and hydrogen-bond distances for WAT1, Gln143, and Trp123. Dotted lines indicate hydrogen bonding and hashed lines indicate SSHBs. **d, e** Tyr34 is observed deprotonated in Mn^3+^SOD and protonated in Mn^2+^SOD. **f, g** A LBHB is seen between His30 and Tyr166, where a proton is transiently shared and is indicated by round dots. The N^δ1^ of His30 is deprotonated when the Mn ion is oxidized and protonated when the Mn ion is reduced. Panel **a** was created from MnSOD X-ray structure (PDB ID 5VF9)^29^, panels **b, d,** and **f** are from the Mn^3+^SOD neutron structure (PDB ID 7KKS)^31^, and panels **c, e,** and **g** are from the Mn^2+^SOD neutron structure (PDB ID 7KKW)^31^. All hydrogen positions were experimentally determined with the exception of solvent molecules in panel **a** that were randomly generated to accentuate the solvent in the active site funnel. **h** Active site overlay of wildtype MnSOD (yellow) and Trp161Phe MnSOD (orange). All distances are in Å.

With neutron protein crystallography of wildtype MnSOD at controlled oxidation states, we observed several changes in protonation and hydrogen bonding in conjunction with changes in the oxidation state of the metal^31^. Neutron crystallography is particularly advantageous for studying PCET mechanisms because neutrons do not alter the electronic state of metals (unlike X-rays), and the neutron scattering of deuterium is on par with carbon, nitrogen, and oxygen^32,33^. The experiment revealed three important pieces of data about the MnSOD mechanism. First, an unusual proton transfer occurred between WAT1 and Gln143 during the Mn^3+^ → Mn^2+^ redox transition where Gln143 transferred an amide proton to a metal-bound ^-^OH molecule, WAT1, to form an amide anion and H_2_O (**Fig 1b, c**). Two short-strong hydrogen bonds (SSHBs) with WAT1 and Trp123 stabilize the amide anion. A SSHB is a type of hydrogen bond that stabilizes catalytic steps and enhances kinetic rates (hashed lines, **Fig 1c**)^34–36^. Second, Tyr34 is deprotonated in the Mn^3+^ redox state and becomes protonated in the Mn^2+^ state (**Fig. 1d, e**). Tyr34 is probably one of the two proton donors needed during *k*_2_ where H_2_O_2_ forms from protonation of substrate^25,27,37,38^. Third, proton transfers occurred between Tyr166 and N^ε2^ of His30 in the form of a low-barrier hydrogen bond (LBHB) that coincide with changes in protonation of N^δ1^ of His30 (round dotted lines, **Fig. 1f, g**). A LBHB is a type of SSHB where the heteroatoms transiently share a proton^39^. Altogether, these studies revealed that electron transfers are coupled with several proton transfer events, and the proximity of the active site metal creates several unusual amino acid pK_a_s that are essential to the mechanism.

Among all SODs, MnSOD has a unique, though poorly-understood, attribute of product inhibition that likely serves to limit the output of H_2_O_2_ within the mitochondrial matrix^30^. H_2_O_2_ is a redox signaling molecule capable of stimulating apoptotic mitochondrial signaling pathways^40,41^ and mitochondrial ‘thiol switches’ to coordinate protein localization and activity^42^. These roles of H_2_O_2_ in mitochondrial redox signaling may explain the evolutionary adaptation of human MnSOD to be more inhibited by H_2_O_2_ compared to its prokaryotic counterparts^26,27^, especially since human abnormalities in H_2_O_2_ steady-state concentrations are hallmarks of disease^43,44^. While the physiological role of MnSOD product inhibition is critical to human health, the biochemical means by which product inhibition is achieved and the identity of the product-inhibited complex is unknown. Structurally revealing the inhibited complex poses benefits for understanding catalysis of MnSOD and, ultimately, the mechanism in which H_2_O_2_ levels are regulated to maintain a beneficial cellular oxidative state.

It has been proposed that product inhibition is initiated from a divalent Mn and O_2_^●-^ to form a complex presumed to be [Mn^3+^-O_2_^2-^] or [Mn^3+^-^-^OOH] (*k*_3_) and is thought to be relieved by at least one protonation to produce Mn^3+^ and H_2_O_2_ (*k*_4_)^26,27^. Note that *k*_2_ and *k*_3_ are competing reactions, both initiated from Mn^2+^, with similar reaction rates (**Table 1**). While the oxidation and protonation states of the inhibited complex (depicted in square parenthesis above and in *k*_3_ and *k*_4_ below) have yet to be determined, the mechanism is attractive as the lack of at least one proton transfer explains why product inhibition occurs, and rate constants are typically calculated with this kinetic mechanism^26,27^. Interestingly, multiple studies have observed that the introduction of H_2_O_2_ with Mn^3+^SOD leads to the formation of the product-inhibited complex that decays to Mn^2+^SOD. However a clear kinetic description of the reaction is lacking^30,45–47^. Mutagenesis studies found the process to be perplexingly correlated with the *k*_2_:*k*_3_ ratio, where a smaller ratio value leads to higher retention of the inhibited complex. This correlation suggests that product inhibition may be more complex than a four-reaction mechanism (i.e. *k*_1_-*k*_4_). Therefore, identifying the inhibited complex is essential for solving the MnSOD product-inhibition mechanism.

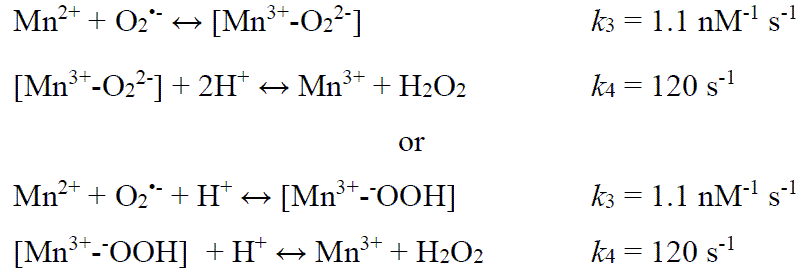

**Table 1.** Individual steady-state rate constants of MnSOD^26^.

| MnSOD | $k_1$ (nM<sup>-1</sup> s<sup>-1</sup>) | $k_2$ (nM<sup>-1</sup> s<sup>-1</sup>) | $k_3$ (nM<sup>-1</sup> s<sup>-1</sup>) | $k_4$ (s<sup>-1</sup>) |
| --- | --- | --- | --- | --- |
| Human Wildtype<sup>a</sup> | 1.5 | 1.1 | 1.1 | 120 |
| Human W161F<sup>b</sup> | 0.3 | <0.01 | 0.46 | 33 |

<sup>a</sup>pH 9.4 and 20 °C from Guan *et al*<sup>48</sup>; <sup>b</sup>pH 9.2 and 25°C from Cabelli *et al*<sup>45</sup>

Current mechanistic insight into human MnSOD product inhibition has relied on kinetic studies of point mutants that alter the competing reaction pathway steps of *k*_2_ and *k*_3_. Of particular interest is the Trp161Phe variant that decreases *k*_2_ by more than two orders of magnitude compared to wildtype, leading to exclusive use of the product-inhibited pathway as *k*_2_ << *k*_3_ (**Table 1**)^26,30,49^. In addition, *k*_4_ is decreased four-fold and indicates higher retention of the product-inhibited complex compared to the wildtype. Of note, the Trp161Phe variant does not remove a titratable amino acid in the hydrogen bond network involved in PCETs. For these reasons, the Trp161Phe variant was selected for experiments to trap and reveal the product-inhibited state.

The Trp161Phe variant slightly distorts the nearby positions of WAT1 and Gln143 suggesting that they are key factors in product inhibition (**Fig. 1h**)^30,48^. Our previous work demonstrated that tight hydrogen bonding between Gln143, WAT1, and Trp123 is present via SSHBs in Mn^2+^SOD (**Fig. 1b**) and may be required for fast PCET catalysis through the uninhibited pathway (i.e. reaction *k*_2_)^31^. Trp161 is immediately adjacent to the WAT1 position and sterically holds the solvent molecule position for interaction with Gln143. The reduction of amino acid size at position 161 from tryptophan to phenylalanine leads to slight movement of WAT1 away from Gln143 (**Fig. 1h**). This lengthening in hydrogen bond distance between WAT1 and Gln143 is hypothesized to have significant catalytic consequences as suggested by the near ablation of the *k*_2_ redox reaction (**Table 1**). Other enzymes that rely on proton transfers for enzymatic activity, such as α-chymotrypsin, substantially increase kinetic rates with SSHBs^35^. Another possible consequence of the Trp161Phe mutation is that WAT1 is more susceptible to displacement from the Mn ion by substrate/product, which would interfere with the back-and-forth proton transfers between Gln143 and WAT1 needed for redox cycling of the Mn ion. Thus, the strategic Trp161Phe variant is advantageous for studying MnSOD catalysis and allows us to determine the significance of the Gln143-WAT1 interaction in product inhibition.

Here, we sought to define the mechanism of human MnSOD product inhibition in terms of PCET catalysis by (1) identifying the mode of product binding to the active site, (2) locating individual proton positions of the product-inhibited state, and (3) determining the electronic configuration of the complex. To pursue these goals, we used neutron crystallography to determine the position of every atom, including hydrogen, at the active site of product-bound Trp161Phe MnSOD, X-ray absorption spectroscopy (XAS) to identify the geometric and electronic structure of the metal and its ligands, and quantum mechanical (QM) chemistry calculations to validate our interpretations of the experimental data. For oxidoreductases, the neutron structures are noteworthy because, to our knowledge, this is one of the first visualizations of a redox center-dioxygen species interaction without radiation-induced artefacts^50^.

## RESULTS AND DISCUSSION

### The structural identity of the product-inhibited complex

To visualize the atomic identity of the product-inhibited complex and the corresponding active site protonation states, a cryocooled neutron structure of perdeuterated Trp161Phe MnSOD soaked with deuterium peroxide (D_2_O_2_) was solved at a resolution of 2.30 Å. We pursued the neutron structure we to visualize a paramagnetic center-oxygen species interaction without radiation-induced perturbations^33^ and identify all the proton positions in the active site. We strategically chose the variant to study product inhibition due to the propensity to enrich and retain the product-inhibited complex at full occupancy without perturbing the hydrogen bond network^26,30,45,46,51^. First, we solved the all-atom structure of the entire enzyme excluding the active site. Then, with these phases we carefully interrogated the active site nuclear density maps for peroxide-derived species that were visually distinct.

The |*F*_o_| – |*F*_c_| nuclear scattering-length density showed that the two subunits of the crystallographic asymmetric unit contained different species bound to the manganese ion opposite His26 (**Fig. 2a**). Differences between the two subunits are often observed for the MnSOD *P*6_1_22 crystal form because the active site of chain *B* is more solvent accessible than the active site of chain *A*^31^. The density of chain *A* has an oval shape that has been previously seen in the unsoaked wildtype isoform and was interpreted as the solvent-bound water molecule, WAT1^31^. For chain *B*, the density is more elongated and in an orientation that is nearly parallel with Asp159. We interpreted the density as a dioxygen species that is singly protonated that has displaced WAT1 to bind the Mn ion (denoted as LIG for ligand, **Fig. 2a**). LIG refined well with full occupancy. This interpretation of the nuclear density is supported by the corresponding X-ray structures (**Supplementary** Fig. 1) and a previous peroxide-soaked *Escherichia coli* wildtype MnSOD structure. However, the dioxygen molecules in these X-ray structures are slightly different in orientations and are at partial occupancies probably due to the detrimental effects of X-ray irradiation^51,52^. At physiological temperatures, optical absorption spectra also suggest a displacement of WAT1 and binding of a dioxygen species, which agrees with our data even though the ligand is cryotrapped^53^. From the nuclear density, we conclude that a singly-protonated dioxygen species replaced a metal-bound solvent molecule to form a five-coordinate complex.

**Fig. 2:**
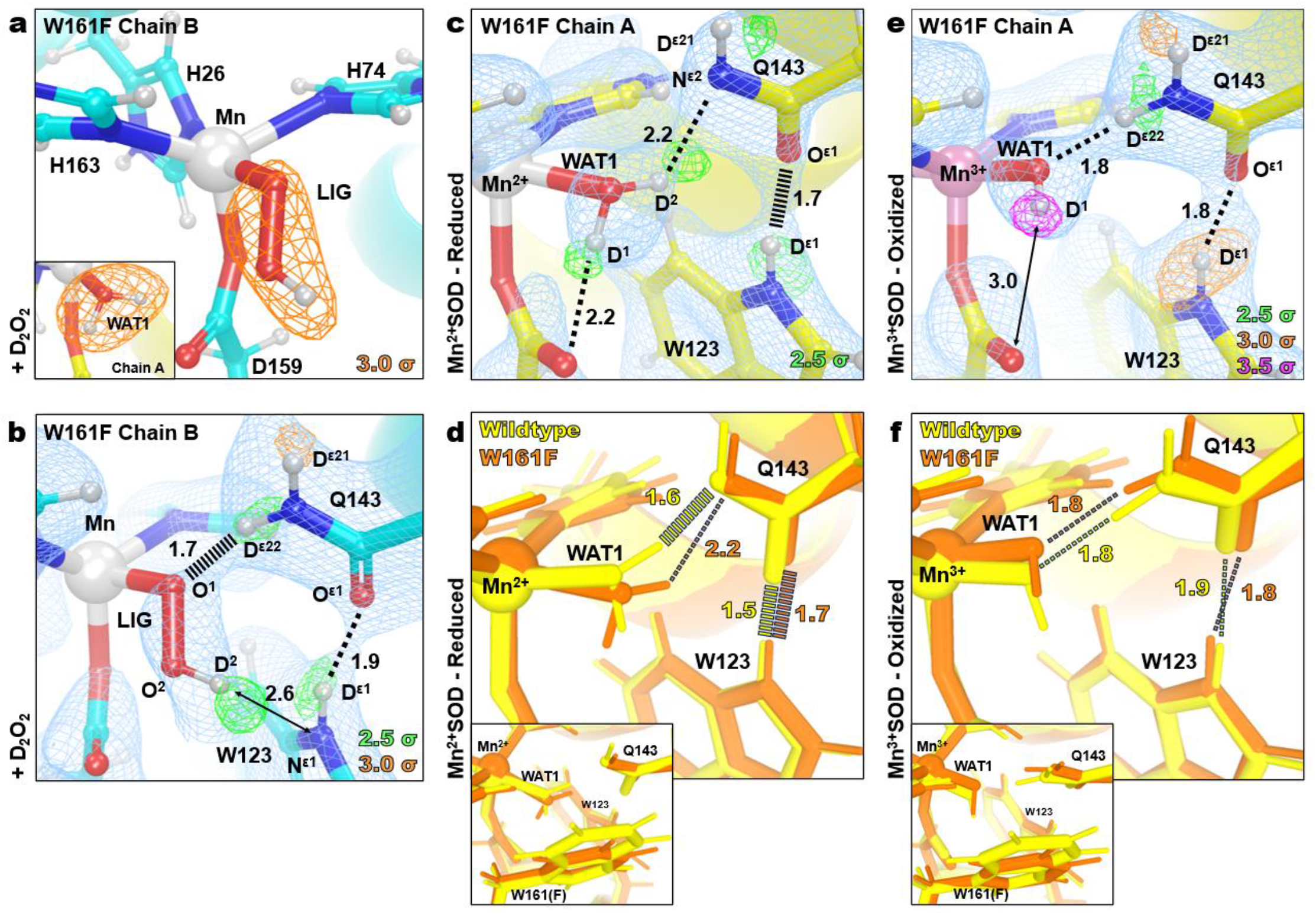
Neutron structures and protonation states at the active site of D_2_O_2_-soaked, reduced, and oxidized Trp161Phe MnSOD. a. D_2_O_2_-soaked Trp161Phe MnSOD for chain *B* with a singly-protonated dioxygen ligand, denoted LIG. Chain A is displayed as an inset to highlight differences in the difference density shapes that led to the identification of LIG in chain B. **b** D_2_O_2_-soaked Trp161Phe MnSOD for chain *B*. **c** Trp161Phe Mn^2+^SOD for chain *A.* **d** Active site overlay of wildtype Mn^2+^SOD and Trp161Phe Mn^2+^SOD demonstrating differences in hydrogen bond strength of the Gln143 amide anion and movement of WAT1. Inset highlights the position of the residue 161 mutation. **e** Trp161Phe Mn^3+^SOD for chain *A*. **f** Active site overlay of wildtype Mn^3+^SOD and Trp161Phe Mn^3+^SOD. Inset highlights the position of the residue 161 mutation. Green, orange, and magenta omit |*F*_o_| – |*F*_c_| difference neutron scattering length density of protons displayed at 2.5σ, 3.0σ, and 3.5σ. respectively. Light blue 2|*F*_o_| – |*F*_c_| density is displayed at 1.0σ. Distances are in Å. Dashed lines indicate typical hydrogen bonds and hashed lines indicate SSHBs that are hydrogen bonds < 1.8 Å.

The protonation states near the Mn ion and the dioxygen ligand that correspond to product inhibition were examined next. The dioxygen species has a single proton (labeled as D^2^, **Fig. 2b**) on O^2^ and points toward the N^ε1^ of Trp123 though it is too far away to be considered a hydrogen bond. Given that the crystal was soaked with D_2_O_2_ before data collection, it can be presumed that the molecule was deprotonated first to form an anion and then displaced WAT1 to bind the positively charged Mn ion. For Gln143, rather than being deprotonated to an amide anion that as seen in wildtype Mn^2+^SOD^31^ and Trp161Phe Mn^2+^SOD (**Fig. 2c, d**), a neutral protonated amide is present. D^ε22^(Gln143) and O^1^(LIG) form a 1.7 Å SSHB, indicating a possible site for proton transfer to occur, especially given that a similar proton transfer was directly observed in wildtype MnSOD (**Fig. 1b, c**). Altogether, a singly-protonated dioxygen ligand bound to Mn that forms a SSHB to a protonated Gln143 has been identified as part of the product-inhibited complex.

The nature of the product-inhibited complex, specifically the coordination of the dioxygen species, has long been under debate^26,27,47,54–56^. One proposed mechanism was that a dioxygen species binds to the sixth ligand site, opposite Asp159. The lack of unfavorable steric interactions and previous DFT energy calculations supported such a complex with a trivalent Mn and a hydroperoxo anion (HOO^-^) as the most probable combination^53–55^. A second proposal was that a peroxyl or hydroperoxyl moiety replaces the position of the solvent ligand as seen in optical absorption spectra and an H_2_O_2_-soaked wildtype *E. coli* MnSOD X-ray crystal structure^52,53^. However, such a five-coordinate inhibited complex in the *E. coli* structure has been interpreted to be less likely as the dioxygen species was partially occupied and introduced steric crowding that leads to higher DFT energies^26,54,55^. X-ray structures are further complicated by the photoreduction induced by X-ray irradiation which may lead to bond distances that are not representative of those under catalytic conditions^33^. Our experimental data from neutron diffraction, which is inherently absent of radiation effects, suggests that product inhibition in human MnSOD is a five-coordinate complex, with a singly-protonated dioxygen ligand displacing the metal-bound solvent molecule. Although there is steric crowding, it is sensible that this prohibits catalysis during the lifetime of the complex as the fast back-and-forth proton transfer between WAT1 and Gln143 (**Fig. 1b, c**) that permits PCET redox cycling is blocked^31^.

### The Gln143-WAT1 SSHB of Mn^2+^SOD limits product inhibition

To look for clues as to what structural features lead to product inhibition, why the Trp161Phe variant is highly product inhibited, and why Trp161 is needed for catalysis, we obtained a cryocooled 2.30 Å resolution neutron structure of Trp161Phe Mn^2+^SOD where the metal was chemically reduced with dithionite (**Fig. 2c**). Dithionite reduces the Mn metal to the divalent oxidation state without entering the active site^31,57^. Indeed, the Mn bond distances of Trp161Phe Mn^2+^SOD are similar to five-coordinate wildtype Mn^2+^SOD (**Supplementary Table 1**). Moreover, the protonation states resemble that of wildtype Mn^2+^SOD, where the bound solvent molecule is D_2_O, not ^-^OD, and Gln143 is deprotonated to an amide anion (**Fig. 1c, 2d**). Deprotonated amino acids are identified when attempts to model and refine a proton result in negative |*F*_o_| – |*F*_c_| difference neutron scattering length density and all the other protons of the amino acid can be placed. Where the Trp161Phe variant and the wildtype contrast is the Gln143-WAT1 interaction. In the wildtype, the Gln143 amide anion forms SSHBs with both WAT1 and Trp123 at distances of 1.6 Å and 1.5 Å, respectively (**Fig. 2d**)^31^. For the variant, there is weak hydrogen bonding between Gln143 and WAT1 with a distance of 2.2 Å while the hydrogen bond of O^ε1^(Gln143) with D^ε1^(Trp123) is weaker compared to wildtype with a distance of 1.7 Å. When comparing the active sites of the wildtype structure versus the Trp161Phe structure, the major difference is the orientation of WAT1 and how closely it interacts with Gln143 (**Fig. 2d**). The oxygen of WAT1 is shifted 0.4 Å away from Gln143 in the variant relative to the wildtype. These observations suggest that for Mn^2+^SOD, the role of Trp161, through its steric repulsion, is to hold Gln143 and WAT1 in a tight hydrogen bonding interaction.

We next investigated the trivalent redox state of MnSOD for structural characteristics that could be assigned to product inhibition. We obtained a room temperature Trp161Phe Mn^3+^SOD neutron structure at 2.30 Å resolution (**Fig. 2e**). Cryocooling and chemical treatment were not pursued due to a limited supply of perdeuterated crystals during beamtime. It is known that purified human MnSOD is ∼90% oxidized^58^. Indeed, the Trp161Phe Mn-ligand bond distances strongly resemble that of the wildtype Mn^3+^ counterpart (**Supplementary Table 2**), and the protonation states are identical, where WAT1 is a ^-^OD molecule and Gln143 is of the protonated amide form (**Fig. 2f**). The variant hydrogen bond distance interactions of Gln143 with WAT1 and Trp123 are identical. Due to the high similarity in structure between Trp161Phe and wildtype in the oxidized state, we conclude that the catalytic consequence of the variant for the Mn^3+^ to Mn^2+^ redox transition is not related to the interactions between the molecules of WAT1, Gln143, and Trp123 but rather pertain to subtle electronic differences or alterations elsewhere in the active site (*vide infra*).

The Trp161Phe variant has historically been puzzling as it profoundly affects PCET catalysis without directly removing a hydrogen bond donor or acceptor^30,45,49^. The diffusion-limited redox transition of Mn^2+^ to Mn^3+^ (*k*_2_, **Table 1**) is nearly ablated while the inhibited complex is enriched (*k*_2_ << *k*_3_, **Table 1**) with a slow disassociation (*k*_4_, **Table 1**)^26,30,49^. From the cryocooled neutron structure of Trp161Phe Mn^2+^SOD (**Fig. 2c**), it is apparent that the Gln143-WAT1 hydrogen bonding interaction is perturbed compared to wildtype Mn^2+^SOD (**Fig. 2d**) and may be the primary cause of the observed kinetic effects. The bulky Trp161 residue that neighbors the Gln143 and WAT1 molecules likely constrains them so that they form SSHBs for proton transfers in the PCET mechanism of MnSOD. We previously established that a strong O(WAT1)-D^2^(WAT1)-N^ε2^(Gln143) interaction, where the proton is covalently but unequally shared, is needed during the Mn^2+^SOD oxidation state for a proton transfer to occur during the Mn^2+^ to Mn^3+^ redox transition^31^. Mutation of glutamine 143 to asparagine similarly ablates catalysis for the Mn^2+^ to Mn^3+^ half-reaction, and a longer WAT1-Gln interaction distance among isoforms of MnSODs and prokaryotic FeSODs correlates with decreased catalytic rates^27,59^. From the neutron structure of Trp161Phe Mn^2+^SOD, we conclude that the role of the Trp161 residue is to promote an interaction between Gln143 and WAT1 amenable to proton transfer the Mn^2+^ to Mn^3+^ redox transition of MnSOD catalysis.

While conserved residue Trp161 has been established to play an integral role in limiting the formation of the product-inhibited complex through kinetic studies, the mechanistic reason has been unknown due to a lack of structural information on the inhibited complex^27,30,45,49^. Our neutron structural data reveals that the inhibited complex is contingent on the displacement of WAT1 by a dioxygen species (**Fig. 2a**) and suggests that the propensity for product inhibition is increased when WAT1 is easier to displace. Indeed, since the WAT1-Gln143 interaction is seen to be destabilized in the Trp161Phe Mn^2+^SOD structure (**Fig. 2d**), WAT1 would be more amenable to replacement by a species that is anionic. Another factor that increases the susceptibility of WAT1 to displacement by an approaching dioxygen species in the Trp161Phe Mn^2+^SOD enzyme is the lower steric repulsion and hydrophobicity of the smaller phenylalanine. The coupling of these two factors potentially explains why the Trp161Phe variant accumulates the inhibited complex and provides an understanding of how human MnSOD regulates product inhibition.

### The electronic identity of the product-inhibited complex

The nature of the product-inhibited complex, specifically the coordination of the dioxygen species and the oxidation state of the Mn ion, has long been under debate^26,27,47,54–56^. Therefore, we pursued Mn K-edge XAS of MnSOD to cross-validate our crystallographic data. K-edge XAS can be divided into two regions: the extended X-ray absorption fine structure (EXAFS) region at higher energies that contains structural information and the X-ray absorption near edge structure (XANES) region that is reflective of the metal oxidation and geometric state. EXAFS of the peroxide-soaked Trp161Phe MnSOD complex was sought to obtain Mn covalent bond distance information for comparison to the neutron structure and QM density functional theory (DFT) calculations (**Table 2**). XANES was similarly pursued to identify the oxidation state and cross-validate the Mn ion’s coordination geometry corresponding to the inhibited complex. X-ray irradiation was carefully monitored so that two subsequent scans of the same spot did not have photoreduction differences, and different positions on the sample were used for each scan.

**Table 2.** Comparison of peroxide-soaked Trp161Phe MnSOD bond lengths from EXAFS fits, neutron structures, and DFT calculations.

| Bond | EXAFS (Å) | Neutron Structure (Å)<sup>b</sup> | DFT (Å)<sup>c</sup> | XANES Fit (Å) |
| --- | --- | --- | --- | --- |
| Mn-N<sup>ε2</sup>(H26) | 2.20 | 2.20 | 2.22 | 2.13 |
| Mn-N<sup>ε2</sup>(H74) | 2.20 | 2.28 | 2.20 | 2.22 |
| Mn-N<sup>ε2</sup>(H163) | 2.20 | 2.17 | 2.17 | 2.16 |
| Mn-O<sup>δ2</sup>(D159) | 2.04 | 2.05 | 2.11 | 2.15 |
| Mn-O<sup>1</sup>(LIG) | 2.04 | 1.94 | 2.04 | 2.04 |
| **Distance** |  |  |  |  |
| Mn-O<sup>2</sup>(LIG)<sup>a</sup> | 2.52 | 2.61 | 2.70 | 2.70 |

<sup>a</sup>The second oxygen of the dioxygen ligand is not directly bound to the Mn ion.
<sup>b</sup>Bond distances correspond to the chain *B* active site of D<sub>2</sub>O<sub>2</sub>-soaked Trp161Phe MnSOD.
<sup>c</sup>DFT calculations utilize a divalent Mn ion with high spin, S=5/2, and a hydroperoxyl anion, HO<sub>2</sub><sup>-</sup>.

We first address the EXAFS structure of peroxide-soaked Trp161Phe MnSOD. The Fourier transform of the raw EXAFS spectra yields the atomic radial distribution around the absorbing Mn ion and is referred to as R space (**Fig. 3a**). The EXAFS spectrum exhibits a peak centered at ∼2.1 Å that is best fit by two sets of scatterers, three nitrogen atoms at 2.20 Å at two oxygen atoms at 2.04 Å (**Supplementary Table 2**). These distances agree with those of Mn ion bond lengths seen in the neutron structure and DFT geometry optimization corresponding with a divalent Mn^2+^ ion (**Table 2**)^31^. The largest contributor to the second peak observed at ∼2.9 are seven carbon atoms that correspond to the C^δ2^ and C^ε1^ of the three Mn-bound histidines and the C^γ^ of the Mn-bound aspartate (**Supplementary Table 2**). For the second oxygen of the dioxygen ligand (O^2^, **Table 2**), the best distance fit for its scattering is at 2.52 Å, which is in slight contrast to the neutron structure and DFT distances of 2.62 Å and 2.70 Å, respectively. This difference may be due to the atom having more degrees of freedom than the Mn-bound atoms. Overall, the structural parameters of the Mn-bound atoms derived from the EXAFS spectrum agree with the crystallographic and QM counterparts.

**Fig. 3:**
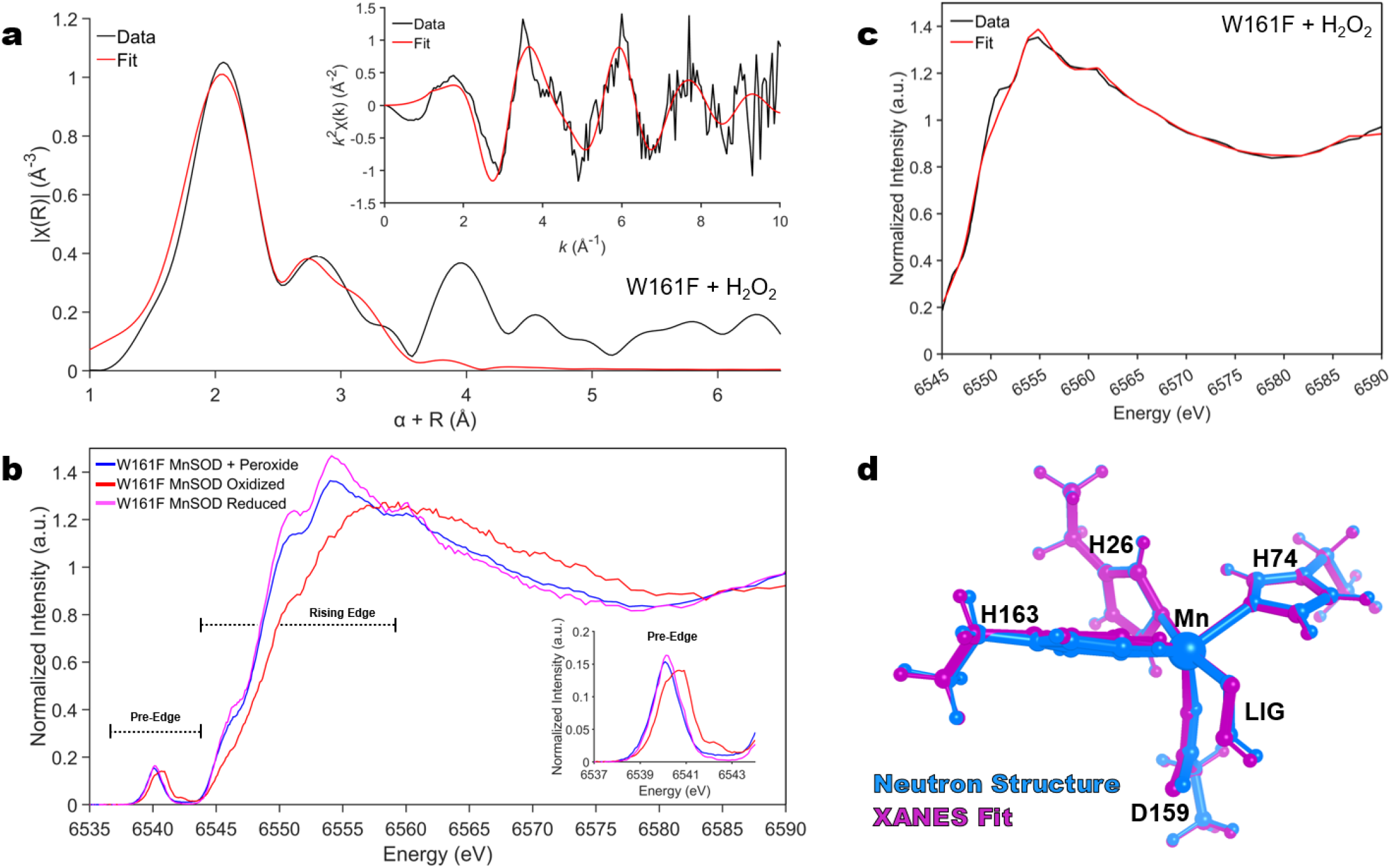
X-ray Absorption Spectroscopy of Trp161Phe MnSOD. a. Fourier transform of Mn K-edge EXAFS data [*k*^2^χ(*k*)] from hydrogen peroxide-soaked Trp161Phe MnSOD with the raw EXAFS spectrum seen in the inset. Due to the scattering phase shift, the distance found by the Fourier Transformation (R) is ∼0.5 Å shorter than the actual distance and a 0.5 Å correction (α) was implemented. The black line represents the experimental data, while the red line is simulated EXAFS spectra from the neutron structure fit to the experimental data. **b** Kα HERFD-XANES of Trp161Phe MnSOD treated with hydrogen peroxide, potassium dichromate, and sodium dithionite to isolate the product-inhibited, Mn^3+^SOD resting, and Mn^2+^SOD resting states, respectively. The inset corresponds to a zoom-in of the pre-edge. **c** Fit of HERFD-XANES spectra of Trp161Phe MnSOD treated with hydrogen peroxide. The black line represents the experimental data, while the red line is simulated XANES spectra from the neutron structure fit to the experimental data. **d** Overlay of the neutron and XANES fit structures that are colored magenta and cyan, respectively.

To obtain the K-edge XANES region, we used the Kα high-energy resolution fluorescence detected absorption (HERFD) method. In K-edge XANES, the oxidation state and coordination geometry can be inferred from the spectra’s rising edge and pre-edge regions. The energy of the rising edge increases as the oxidation state of the metal increases while staying in a high spin state. When comparing the rising edges in the XANES region between oxidized and reduced Trp161Phe MnSOD, a shift to lower energy is observed that indicates a one-electron difference of oxidation state and coincides with Mn^3+^SOD and Mn^2+^SOD resting forms, respectively (**Fig. 3b**)^60^. With the presence of hydrogen peroxide, the rising edge is most like the reduced counterpart, indicating that the Mn metal is in a divalent oxidation state. The feature around 6554 eV of peroxide-soaked Trp161Phe MnSOD is of lower intensity than the reduced form while a higher intensity is observed for peroxide-soaked Trp161Phe MnSOD at ∼6570 eV. These differences, in conjunction with noticeable variation in the overall shape resonance (6540-6590 eV) between peroxide-soaked Trp161Phe MnSOD and reduced Trp161Phe MnSOD, reflect an alteration in structure that is presumably caused by replacement of the solvent molecule with the dioxygen species in peroxide-soaked Trp161Phe MnSOD. The HERFD-XANES spectra for wildtype MnSOD demonstrate the same trends; however, the dioxygen-bound chemical species from peroxide soaking is not as well isolated due to the lesser extent of product inhibition that the wildtype form has compared to the Trp161Phe variant (**Supplementary** Fig. 2). For the pre-edge region, the intensity (i.e., area of the pre-edge feature) corresponds to metal 1s to 3d transitions that gain intensity through 4p character mixed into the 3d orbitals as a result of symmetry distortions and a loss of (local) inversion symmetry, with the mixing mediated by metal-ligand covalency^61–63^. The influence of symmetry is of particular note as a six-coordinate Mn complex is expected to have different pre-edge intensity compared to a five-coordinate complex. For Trp161Phe with peroxide treatment, the pre-edge intensity is comparable to both the oxidized and reduced and indicates the coordination number is the same between the three chemical species (**Fig. 3b**). Like the rising edge, the pre-edge also shifts in energy depending on the oxidation state of the metal. A higher energy pre-edge for oxidized Trp161Phe MnSOD coincides with a Mn^3+^ ion, while a lower energy pre-edge for reduced and peroxide-soaked Trp161Phe MnSOD coincides with a Mn^2+^ ion. From the HERFD-XANES data, we can conclude that the introduction of hydrogen peroxide leads to a divalent active site Mn metal that is five-coordinate, though it differs in structure compared to the reduced resting state.

Next, we performed structural refinement using the shape resonance of the HERFD-XANES spectra between 6545 and 6590 eV using a combined DFT and machine learning approach previously developed^64,65^. With this method, systematic deformations of an input neutron structure were applied, and the corresponding XANES spectra of each deformation were simulated. The simulated spectra of the deformed structures then served as a training set for assigning a structure-to-spectra relationship for structural refinement. For the peroxide-soaked Trp161Phe complex, the simulated spectra of the best-fit structure captures nearly all features and relative intensities of the experimental spectra (**Fig. 3c, d**) except for the 6550 eV feature that is found by the fit but underestimated. These trends are also the case for oxidized and reduced counterparts for the variant and wildtype (**Supplementary** Fig. 3). Likewise, the Mn-bond distances of the XANES structural refinement are in agreement with the structures from EXAFS, neutron diffraction, and DFT (**Table 2 and Supplementary Table 3**), with the largest difference of bond length for peroxide-soaked Trp161Phe MnSOD being 0.1 Å. Overall, the structural information of the inhibited complex extracted from the shape resonance of the HERFD-XANES spectra coincides with the structure derived from other methods and gives further confidence in the identity of the complex.

Clear evidence for the oxidation state of the Mn during product inhibition has previously been challenging to obtain due to ambiguities in optical spectra, large zero-field splitting values for the Mn using conventional electron paramagnetic resonance (EPR), and efforts to capture the inhibited complex with the wildtype enzyme leading to a mixture of species^26,27,45–47,59^. Historically, the product-inhibited oxidation state has generally been accepted to be Mn^3+^ as (1) this is the oxidation state observed following decay of the product-inhibited complex, (2) oxidative addition of dioxygen species to low-valent metal have been described, and (3) disassociation of product inhibition exhibits a solvent-hydrogen isotope effect, which has been interpreted as relief of the complex being proton transfer, and not electron transfer, dependent^26,27,45–47^. However, the HERFD data of the Mn ion in both the rising edge and pre-edge regions with the introduction of hydrogen peroxide to the Trp161Phe variant (**Fig. 3b**), and the metal bond distances observed by neutron diffraction and EXAFs resemble those of a divalent Mn ion (**Table 2**). Previous wildtype and Trp161Phe studies that coupled kinetics and visible spectra demonstrate that the inhibited complex lacks the characteristic 480 nm absorbance of the five-coordinate trivalent Mn oxidation state, although they were unable to rule out a six-coordinate Mn^3+^ complex^30,45,46^. These past observations can be reconciled with our new data that indicate a five-coordinate Mn^2+^ complex.

Thus far, we have not addressed the electronic identity of the singly-protonated dioxygen ligand. Is it a protonated superoxide (HO_2_^•^) or hydroperoxyl anion (HO_2_^-^)? Given that H_2_O_2_ soaking leads to one-electron reduction of the metal^30,45,47^, one possible source of the electron is from H_2_O_2_ forming HO_2_^•^ in conjunction with a proton transfer. A second possibility is after the reduction of Mn^3+^SOD by H_2_O_2_ leading to Mn^2+^SOD, another H_2_O_2_ molecule is deprotonated and binds the metal without electron transfer. At this time, we cannot experimentally differentiate between HO_2_^•^ and HO_2_^-^. Our DFT calculations yield Mn ligand bond distances that most closely resemble our experimental data when the identity of the dioxygen species is HO_2_^-^ (**Table 2**). Compared to HO_2_^•^, HO_2_^-^ is a more attractive candidate to displace the WAT1 ligand given its anionic charge. HO_2_^-^ is also more chemically likely to lead to a stable product-inhibited complex due to its non-radical status. Furthermore, this suggestion does not discount past solvent-hydrogen isotope effect studies that observe relief of the product-inhibited complex as proton-transfer dependent, as protonation of HO_2_^-^ to H_2_O_2_ would cause disassociation of the dioxygen species^30^. While we cannot definitively ascribe an electronic identity to the dioxygen species, our quantum chemistry calculations (**Table 2**) support the interpretation of an HO_2_^-^ molecule.

### Determination of the molecular orbitals that define redox reactivity

We next investigated the molecular orbitals that determine the enzyme’s reactivity with DFT calculations and analysis of the pre-edge HERFD-XANES spectra. First, we performed DFT calculations to describe the ground state of wildtype Mn^3+^SOD, wildtype Mn^2+^SOD, and Trp161Phe Mn^2+^SOD bound to HO_2_^-^. These three complexes were selected as they reflect the most physiologically relevant complexes that had experimental data. Unambiguous structural data of wildtype MnSOD bound to HO_2_^-^ is unavailable, so the Trp161Phe variant was used instead. Wildtype MnSOD is known to be high spin S=2 and S=5/2 for the trivalent and divalent states, respectively^66^.

First, we focus on ground-state DFT calculations for Mn^3+^SOD as its unoccupied metal 3d orbitals largely determine the redox reactivity of the enzyme. MnSOD is best described by a distorted local *C*_3*v*_ symmetry where the z-axis is along the Mn-oxo bond of the Mn-WAT1 interaction (**Fig. 4a**). High spin (S=2) Mn^3+^ has one unoccupied α-spin orbital and five unoccupied β-spin orbitals. The lowest unoccupied molecular orbital (LUMO) is composed predominately of the 3d_z_^2^ α orbital due to strong spin polarization from the partially occupied α-manifold and interacts strongly with ^-^OH. The LUMO is composed of 57% d character while the ^-^OH contribution to the orbital is 17% indicating that the ^-^OH molecule is a contributor towards reactivity of the orbital. The e_π_ (xz/yz) orbitals of the β-manifold have strong d character of 81%, though are tilted along the axis suggesting mixing with other d orbitals. The similarly tilted e_σ_ (xy/x^2^-y^2^) β orbitals have 74% d character and bond strongly with the amino acid ligands leading to 1.5% mixing of the Mn 4p orbitals with the e_σ_ orbitals. Due to the 4p mixing, the e_σ_ orbitals are expected to lead to significant electric dipole character even with these small percentages of mixing^61,62,67^. Lastly, the 3d_z_^2^ β orbital is strongly σ-interacting with ligands along the z-axis. The 2.1 eV splitting between the z^2^ α and β orbitals is explained by spin polarization differences between the α– and β-manifolds reflected by a partially occupied α-manifold and an entirely unoccupied β-manifold. From the ground-state DFT calculations of Mn^3+^SOD, the 3d_z_^2^ α orbital is expected to be the orbital participating in enzymatic redox reactions, and the ^-^OH molecule contributes towards reactivity.

**Fig. 4:**
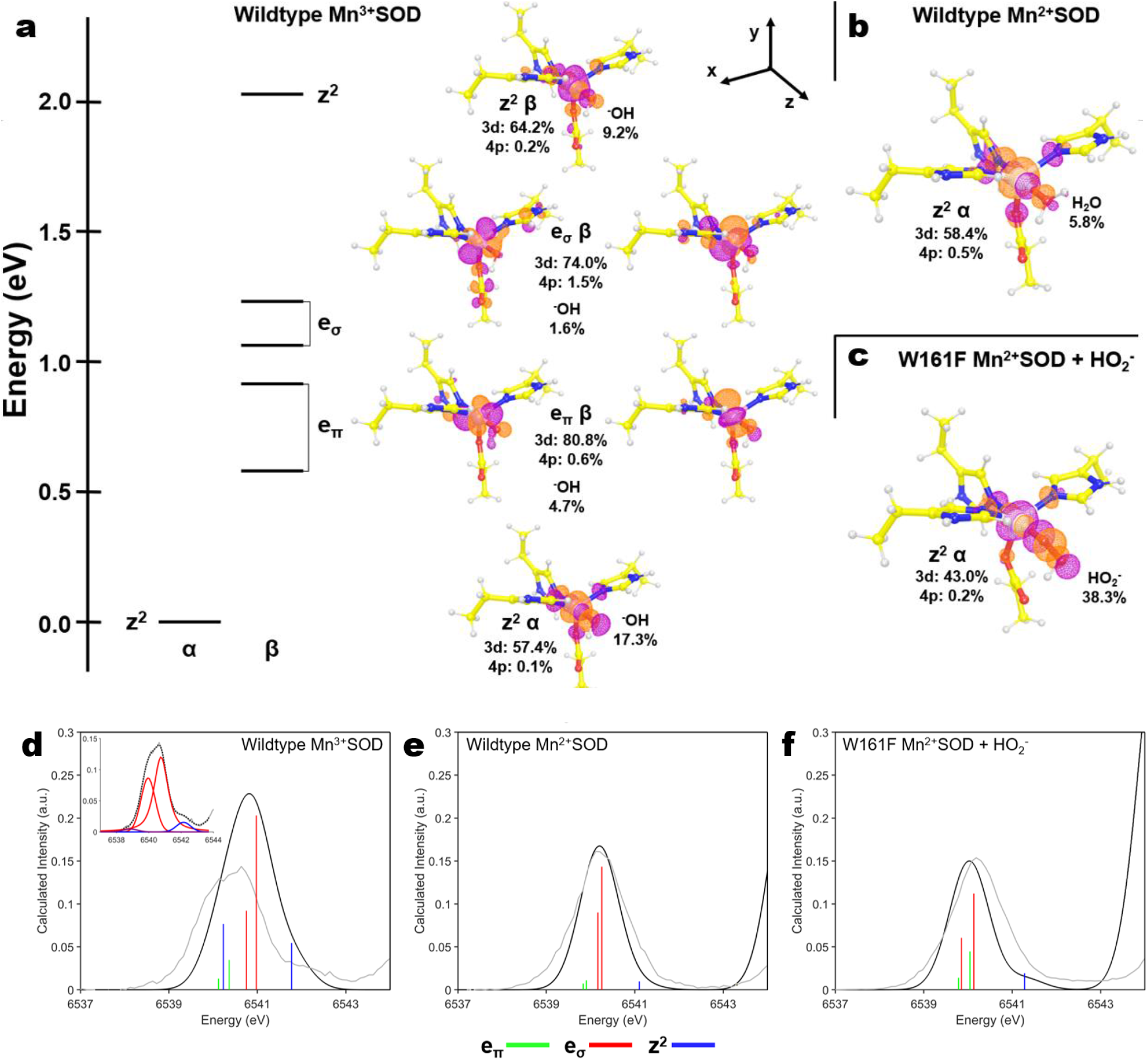
DFT simulations of wildtype Mn^3+^SOD, Mn^2+^SOD, and Trp161Phe Mn^2+^SOD bound to HO_2_^-^. a. Energy diagram of the unoccupied valence orbitals of Mn^3+^SOD. The energy axis is shifted to set the z^2^ α orbital at zero. The isosurface plots of the unoccupied orbitals are contoured at 0.05 au. Percentages of orbital character are derived from Löwdin population analysis where the *C*_3v_ symmetry-related xz and yz orbitals were averaged for the e_π_ values, and the xy and x^2^-y^2^ orbitals were averaged for the e_σ_ values. **b** The HOMO of Mn^2+^SOD. **c** The HOMO of Trp161Phe Mn^2+^SOD bound to HO_2_^-^. **d-f** TDDFT simulated spectra where the simulated spectrum is colored black, and the corresponding experimental HERFD-XANES spectrum is colored gray. The green, red, and blue vertical lines correspond to e_π_, e_σ_, and z^2^ transitions, respectively. Simulated intensities were uniformly scaled to experimental intensities, and the energy axis was shifted. Due to a misestimation of the Mn^3+^SOD pre-edge spectra from the TDDFT simulation (see text), a pseudo-Voight peak fit was performed with identical color-coding of the transitions and is included as an inset for panel **d**. Unclear peak identities are colored purple.

Ground-state DFT calculations were performed to analyze the highest occupied molecular orbitals (HOMOs) of Mn^2+^SOD and Trp161Phe Mn^2+^SOD bound to HO_2_^-^. Here, we focused on the HOMOs as these are the orbitals participating in catalytic activity. In both Mn^2+^ complexes, the 3d ^2^ α orbital is the HOMO and interacts with the H_2_O or HO_2_^-^ ligand along the z-axis (**Fig. 4b, c**). For wildtype Mn^2+^SOD the d character of 58% strongly resembles that of the unoccupied 3d ^2^ α orbital of Mn^3+^SOD with 57% d character (**Fig. 4a**). Given the protonation of the solvent-ligand in Mn^2+^SOD (**Fig. 4b**), the 2p character from the solvent contributes less density to the 3d ^2^ α orbital compared to that of Mn^3+^SOD. In the case of Trp161Phe Mn^2+^SOD bound by HO_2_^-^, the HO_2_^-^ contributes a large portion of density to the orbital, with Mn 3d and HO_2_^-^ contributions of 43% and 38%, respectively (**Fig. 4c**). The density localized on the HO_2_^-^ is reflective of the 2p π-antibonding orbital. Since the z^2^ α orbital performs redox reactions, it is sensible that any oxo or dioxo species involved in catalysis would bind in an orientation favoring interaction along the z-axis as it leads to the greatest orbital overlap. Thus, analysis of the HOMOs potentially explains why HO_2_^-^ prefers to displace the metal-bound solvent rather than binding opposite the aspartate residue.

Next, we calculated the Mn K-pre-edge XAS for the three complexes using time-dependent density functional theory (TDDFT), an excited-state simulation, to assign the metal 3d orbital contributions to the experimental HERFD-XANES spectra of the pre-edge. The simulated spectra and corresponding experimental spectra are plotted in **Fig. 4d-f**. The vertical stick heights correspond to the intensity of the 1s to 3d transitions with the sum of quadrupole and dipole contributions. Overall, the spectral shapes of the divalent complexes (**Fig. 4e, f**) are well reproduced by the simulation. On the other hand, for the trivalent complex (**Fig. 4d**) the splitting of the orbital transitions is underestimated while the intensity is overestimated.

The TDDFT simulation of wildtype Mn^3+^SOD (**Fig. 4d**) identifies the e_σ_ transitions as the primary contributors of intensity, although it has a misestimation of the overall spectral intensity and shape. Significant dipole intensity is found in the e_σ_ and z^2^ transitions. The e_σ_ orbitals bear the most dipole character which is reflective of the higher 4p mixing seen in the ground-state DFT calculations (**Fig. 4a**). The valence 3d orbital splitting of the simulation is underestimated and the contribution of the transitions to the spectra is better represented by pseudo-Voigt peak fitting of the experimental data that suggests an increased splitting energy (inset, **Fig. 4d**). Four peaks were required to fit the spectrum and attempts to fit five or more peaks did not lead to a better solution. The two larger peaks found in the fit potentially stem from splitting of the e_σ_ orbitals (xy/x^2^-y^2^) due to the distorted *C*_3v_ symmetry where one of the orbitals may have more overlap with those of ligands. The higher energy peak found in the fit at ∼6542.5 eV is best identified as the z^2^ β transition supported by both ground and excited-state DFT. A fourth peak is capable of being fit as a small shoulder at ∼ 6539 eV though the orbital contribution to the peak is unclear as it could be of e_π_ or z^2^ α character (or both). The e_π_ or z^2^ α orbitals are probably underneath the stronger e_σ_ orbitals. For the TDDFT simulation, the overestimation of the transition intensities and underestimation of the valence 3d orbital splitting has been previously noted for other high spin

S=2 d^4^ complexes with similar symmetry^67,68^. In brief, the Mn 1s core hole exchange interaction with the valence 3d orbitals is overestimated by DFT, and its effects in the spectra are most prominent when there are both α and β transitions. This overestimation explains why the misestimations are less pronounced in the divalent S=5/2 d^5^ complexes (**Fig. 4e, f**) as they contain only β transitions. Nevertheless, the TDDFT simulation suggests that the e_σ_ transitions are the predominant contributors to pre-edge spectral intensity for wildtype Mn^3+^SOD.

For Mn^2+^SOD, the assignment of the metal 3d orbital contributions to the experimental spectra is more straightforward. The TDDFT simulation of wildtype Mn^2+^SOD (**Fig. 4e**) suggests the e_σ_ orbitals have the most dipole character from 4p mixing and thus predominantly contribute to the intensity. The e_π_ orbital transitions are mostly of quadrupole character, leading to a weak low-energy tail. Note that the e_π_ orbitals are allowed to mix with the 4p orbitals though due to π-bonding, the effect is small^63^. The z^2^ transition consists of low dipole intensity leading to a high energy tail. The trend of transition energy (e_π_, e_σ_, z^2^) mirrors that of Mn^3+^SOD (**Fig. 4d**) and suggests the order of the metal 3d molecular orbital energies are maintained between the two resting redox states. Overall, the TDDFT simulation of wildtype Mn^2+^SOD matches the experimental data.

For the TDDFT simulation of Trp161Phe Mn^2+^SOD bound to HO_2_^-^ (**Fig. 4f**), the overall spectra shape agrees well with that of the experimental data. Here, all transitions bear significant dipole intensity, the e_π_ and e_σ_ transitions overlap in energy, and the z^2^ transition contributes a tail to the spectra. The observations coincide with further distortion of *C*_3v_-symmetry introduced from the binding of the HO_2_^-^ molecule, causing the metal 3d orbitals to mix further. This mixing distributes the dipole character over the five unoccupied β orbitals, leading to an overlap of the e_π_ and e_σ_ sets of orbitals. Importantly, the mixing permits distinction of a dioxygen species-bound complex against that of a solvent molecule-bound complex seen in Mn^3+^ and Mn^2+^ resting states (**Fig. 4d, e**). Overall, the TDDFT simulation of Trp161Phe Mn^2+^SOD bound to HO_2_^-^ agrees with the experimental data.

The ground-state and excited-state DFT simulations help explain the catalytic mechanism of MnSOD. The S=2 d^4^ configuration of Mn^3+^SOD and the distorted trigonal ligand field leads to spin polarization of the 3d_z_^2^ α orbital and its large splitting with the 3d_z_^2^ β orbital (**Fig. 4a**). This splitting is reflected in the experimental pre-edge spectra though is underestimated in the TDDFT simulations as documented by studies of similar complexes (**Fig. 4d**)^67,68^. Importantly, the splitting leads to the 3d_z_^2^ α becoming the LUMO that is the putative electron acceptor orbital for reaction with O_2_^•−^. The 3d_z_^2^ α orbital mixes strongly with antibonding molecular orbitals of the metal-bound ligands. In particular, the ^-^OH molecule bound along the z-axis contributes a relatively large portion of density towards the LUMO to suggest that it is a determinant in redox activity. For Mn^2+^SOD that has an S=5/2 d^5^ configuration, ground-state DFT calculations (**Fig. 4b**) indicate that the 3d_z_^2^ α orbital is the HOMO, the putative electron donor orbital for reaction with O_2_^•−^, while the TDDFT simulation (**Fig. 4e**) matches the HERFD-XANES spectra suggesting that our calculations are recapturing the experimental data for the complex. For the Trp161Phe Mn^2+^SOD complex bound to HO_2_^-^, the HOMO is again the 3d_z_^2^ α orbital though the 2p π-antibonding character of HO_2_^-^ contributes a large portion of density to the orbital (**Fig. 4c**). As previously hypothesized by Jackson and colleagues^69^, the interaction demonstrates why displacement of the metal-bound solvent and binding along the z-axis is preferred for a dioxygen species since the dioxygen π-antibonding orbitals overlap with the Mn 3d ^2^ α orbital. Additionally, the experimental and TDDFT simulations of the XAS pre-edge (**Fig. 4f**) permit spectral discrimination of a dioxygen-bound Mn^2+^SOD complex against a resting state Mn^2+^SOD complex. Altogether, the DFT simulations indicate that (1) the z^2^ α orbital is the orbital performing redox reactions, (2) reactivity is influenced by the species bound along the z-axis of the Mn, and (3) displacement of the solvent molecule for dioxygen species binding is chemically preferred due to orbital overlaps.

### Unexpected second-sphere hydrogen peroxide binding site

We previously demonstrated that residues Tyr166, His30, and Tyr34 all have unusual pK_a_s and undergo changes in protonation states during the MnSOD PCET cycle (**Fig. 1**)^31^. Specifically, the active site gains two protons during the Mn^3+^ → Mn^2+^ redox transition. One proton is acquired by Tyr34 and the other by His30. Tyr166 modulates His30 with a LBHB. These protons are lost during the Mn^2+^ → Mn^3+^ transition and this led us, at the time, to hypothesize that Tyr34 and His30 bind and protonate O_2_^2-^ to H_2_O_2_, especially since the gap between these two residues is the only solvent-accessible entrance/exit of the active site. When investigating the active site of the D_2_O_2_-soaked structure at chain *B*, a strong omit |*F*_o_| – |*F*_c_| difference density peak is seen between His30 and Tyr34 with an oblong shape that is longer than a water molecule (orange density, **Fig. 5a**). This is the same active site where the dioxygen ligand is bound to the Mn^2+^ ion (**Fig. 2a**). We interpret the omit nuclear density as D_2_O_2_ (denoted as PEO, **Fig. 3a**) given (1) the crystal was soaked with D_2_O_2_ before data collection, (2) modeling and refining a D_2_O (water) molecule leads to unoccupied residual density, and (3) the density has two protruding features suggestive of two protons. The locations of the D_2_O_2_ protons are further verified by their nuclear density peaks (green density of PEO, **Fig. 5a**), and it is expected that the density of a proton engaging in strong hydrogen bonding, such as D^1^(PEO), to be more localized and defined compared to a proton that is rotatable and lacking hydrogen bond acceptors, such as D^2^(PEO). Indeed, the D^1^(PEO) proton forms a 1.4 Å SSHB with a deprotonated Tyr34. Furthermore, the D_2_O_2_ molecule forms a second 1.4 Å SSHB with His30, between atoms D^δ1^(His30) and O^1^(PEO). The proton between O^η^(Tyr166) and N^ε2^(His30) that we previously observed participating in a LBHB for the resting Mn^2+^SOD structure (**Fig. 1g**) is instead fully localized onto O^η^(Tyr166) and participating in a 1.6 Å SSHB with N^ε2^(His30). The experimental data suggest a strong network of hydrogen bonds is present at physiological pH involving Tyr34-PEO-His30-Tyr166 supporting our hypothesis that Tyr34 and His30 bind and protonate O ^2-^ to H_2_O_2_

**Fig. 5:**
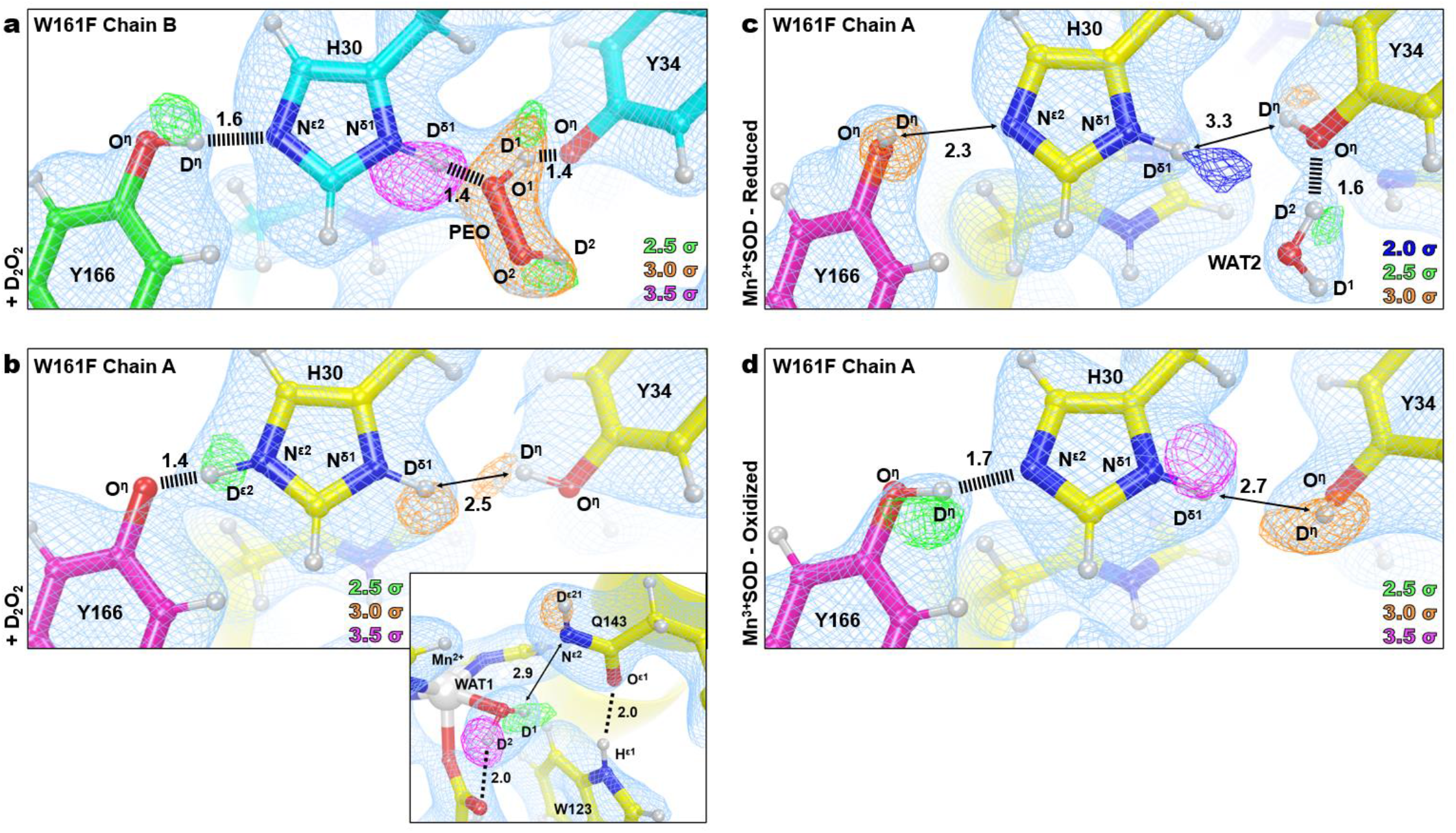
Neutron structures and protonation states of second sphere active site residues in D_2_O_2_-soaked, reduced, and oxidized Trp161Phe MnSOD. a. D_2_O_2_-soaked Trp161Phe MnSOD at the active site of chain *B*. **b** D_2_O_2_-spaked Trp161Phe MnSOD at the active site of chain *A*. Inset highlights the structure near the divalent Mn ion. Chain *B* is more accessible to solvent than chain *A* and helps explain differences in ligand binding. **c** Divalent resting state of Trp161Phe MnSOD at the active site of chain *A*. **d** Trivalent resting state of Trp161Phe MnSOD at the active site of chain *A.* Blue, green, orange, and magenta omit |*F*_o_| – |*F*_c_| difference neutron scattering length density of protons are displayed at 2.0 σ, 2.5σ, 3.0σ, and 3.5σ, respectively. Light blue 2|*F*_o_| – |*F*_c_| density is displayed at 1.0σ. Distances are in Å. Dashed lines indicate typical hydrogen bonds, and hashed lines indicate SSHBs, hydrogen bonds < 1.8 Å. For the resting state structures, only one active site is shown due to high structural similarities, see **Supplementary** Fig. 4 for the other active site.

The active site of chain *A* for the D_2_O_2_-soaked Trp161Phe MnSOD, which does not have binding of a dioxygen species in either the first or second spheres, was next investigated for insight on the effect of dioxygen species binding on protonation (**Fig. 5b**). The active site has a divalent Mn^2+^ ion with Mn bond distances and protonation states reflecting that of Trp161Phe and wildtype Mn^2+^SOD (**Supplementary Table 1**). Without treatment of redox reagents or D_2_O_2_/H_2_O_2_, human MnSOD is ∼90% trivalent^58^ (**Fig. 2e and Supplementary** Fig. 2) and suggests a peroxide molecule performed one-electron reduction with the metal, and the resulting species vacated the active site before flash-freezing. Here, it is also evident that the variant is destabilizing the Gln143-WAT1 interaction as the two molecules are not in a hydrogen bond (**Fig. 5b**, inset). The omit |*F*_o_| – |*F*_c_| difference density suggests that Tyr34 is protonated (**Fig. 5b**) and contrasts with the Tyr34 of chain B, suggesting that Tyr34 gains a proton from D_2_O_2_. For His30 of chain A, omit |*F*_o_| – |*F*_c_| difference density is observed for protons on both nitrogen atoms at contours of 2.5σ or higher, indicating His30 is positively charged and Tyr166 is negatively charged. The D^ε2^(His30) proton forms a 1.4 Å SSHB with O^η^(Tyr166) and also contrasts with chain B in that the proton is localized on His30 rather than Tyr166. Given that the proton between N^ε2^(His30) and O^η^(Tyr34) is seen altering its position, a back-and-forth proton transfer may occur between the heteroatoms depending on the ligand environment, such as the binding of a dioxygen species. The observation also reinforces the model originated from the room temperature wildtype neutron structures where a proton is either transiently shared between His30 and Tyr166, or a back-and-forth proton shuffle occurs during the catalytic cycle^31^. Overall, the neutron structure of D_2_O_2_-soaked Trp161Phe MnSOD at the active site of chain *A*, in conjunction with the active site of chain *B*, provides insight into the role of Tyr34, His30, and Tyr166 during product binding.

We next examined the neutron structures of resting state Trp161Phe Mn^2+^SOD and Trp161Phe Mn^3+^SOD for contextual clues of the product inhibition mechanism and overall catalysis. Both chains of each structure have high structural similarity (**Supplementary** Fig. 4). For Trp161Phe Mn^2+^SOD, the protonation states of Tyr34, His30, and Tyr166 resemble those of the wildtype Mn^2+^SOD though there are notable structural features. Tyr34 engages in a 1.6 Å SSHB with WAT2 where O^η^(Tyr34) acts as a hydrogen-bond acceptor (**Fig. 5c**). From our past work, it is apparent that Tyr34 forms SSHBs with water molecules regardless of its protonation state and theoxidation state of the Mn ion and supports the notion that Tyr34 is central to proton turnover during catalysis^31^. Indeed, its mutation predominately affects the Mn^2+^ to Mn^3+^ redox transition of the enzyme that has a solvent-exchangeable proton as the rate-limiting step of the half-reaction^26,27,49^. Besides Tyr34, the other notable structural feature is the hydroxyl proton of Tyr166 that points away from His30 (**Fig. 5c**). It is unclear whether the orientation of the proton has catalytic significance, is somehow related to the Trp161Phe variant, or is a consequence of cryocooling. Regardless, the Trp161Phe Mn^2+^SOD and the D_2_O_2_-bound neutron structures (**Fig. 5a**), provide evidence that Tyr34 is involved in SSHBs with both D_2_O_2_ and water molecules and supports the interpretation that Tyr34 gives and takes protons during catalysis.

Surprisingly, Tyr34 and His30 are both protonated in the neutron structure of Trp161Phe Mn^3+^SOD, which is the opposite of wildtype Mn^3+^SOD protonation states (**Fig. 1d, f**). Strong omit |*F*_o_| – |*F*_c_| difference density is observed for D^η^(Tyr34), D^δ1^(His30), and D^η^(Tyr166) that forms a 1.7 Å SSHB with N^ε2^(His30) (**Fig. 5d**). A potential explanation for the contrasting protonation states compared to wildtype Mn^3+^SOD are alterations of residue pK_a_s due to small changes in the active site. Past studies across different isoforms of MnSOD with conserved active sites have established that subtle changes in hydrogen bonding and residue orientation lead to significant changes in catalysis, mainly due to shifts of the net electrostatic vectors^26,27,29,56,70^. Other changes in the Trp161Phe variant include the previously discussed Gln143 and WAT1 molecules (**Fig. 2f**) and a slight 1.8 Å movement of Tyr34 toward His30 (**Supplementary** Fig. 5). Our neutron diffraction data of Trp161Phe Mn^3+^SOD suggest that subtle movement of active site residues affect the pK_a_ of residues involved in proton transfer for PCET catalysis.

Overall, our Trp161Phe MnSOD neutron structures indicate that D_2_O_2_ hydrogen bonds strongly to His30 and Tyr34 and confirms that residues Tyr34, His30, and Tyr166 undergo protonation changes between catalytic states. The gateway binding site of D_2_O_2_ between His30 and Tyr34 was predicted though, to our knowledge, the neutron structure is the first experimental evidence for this binding^26,27,31,38,56,71^. The gateway binding site has been thought to be where an O_2_^2-^ or HO_2_^-^ species gain protons during the Mn^2+^ to Mn^3+^ redox transition (*k*_2_). However, our neutron structure of D_2_O_2_-soaked Trp161Phe MnSOD brings the possibility that it may also play a role during product inhibition, with three possible interpretations of the data. First, D_2_O_2_ binding occludes the only entrance to the active site and may potentially block substrate from entering since the Tyr166-His30-PEO-Tyr34 network is composed of several SSHBs (**Fig. 5a**). Second, the life of the network is short-lived and a PEO → Tyr34 proton transfer would create a HO_2_^-^ with high electrostatic affinity towards the Mn ion. Third, a HO_2_^-^ molecule already bound to Mn^2+^ may stabilize H_2_O_2_ species binding between His30 and Tyr34 and block the product from exiting. Altogether, the neutron structures provide considerable insight into MnSOD catalysis by revealing both binding orientations of dioxygen species and protonation states.

### Summary of active site configurations solved

The neutron structures, XAS data, and simulations of D_2_O_2_-soaked, reduced, and oxidized Trp161Phe MnSOD present four primary configurations of active sites. For chain *B* of the D_2_O_2_-soaked structure, two dioxygen species are found within the active site (**Fig. 6a**). One of the dioxygen species is singly protonated and bound directly to the metal, replacing the position of WAT1, and forming a 1.7 Å SSHB with Gln143 that is in the canonical, protonated amide form (**Fig. 2a**). Our XAS measurements and DFT simulations support the interpretation that the Mn ion is divalent and the singly-protonated dioxygen species is HO_2_^-^ (**Fig. 3**, **Fig. 4**, **Table 2**). Between an ionic Tyr34 and neutral His30 of the same active site, D_2_O_2_ is observed binding tightly by forming 1.4 Å SSHBs with gateway residues, and Tyr166 forms a 1.6 Å SSHB with His30 (**Fig. 5a**). For chain *A* of the same D_2_O_2_-soaked neutron structure, the active site is reduced but a dioxygen ligand is not present (**Fig. 6b**). Since peroxide is established to reduce the Mn ion (**Fig. 3b**)^30,45,46^, we presume that the active site metal underwent reduction with a peroxide molecule and the resulting dioxygen species left before cryocooling. Gln143 is in the unusual, unprotonated amide anion form where the dominant resonance form is a double bond between N^ε2^ and C^δ^ with the O^ε1^ atom bearing the negative charge. Tyr34 is observed in the neutral state while His30 is protonated on both nitrogen atoms and is positively charged. Tyr166 is ionized. The interaction between His30 and Tyr166 is intimate as they form a 1.4 Å SSHB suggestive of a proton transfer site (**Fig. 5b**). For the Trp161Phe reduced resting state, both chains have the same structure, and WAT1 is engaging in a 2.2 Å hydrogen bond with an amide anion form of Gln143 (**Fig. 6c**). Due to its higher anionic character, O^ε1^(Gln143) forms a 1.7 Å SSHB with H^ε1^(Trp123). Tyr34 is protonated and forms a SSHB with WAT2 while His30 and Tyr166 do not engage in hydrogen bonds. The Trp161Phe oxidized resting state also has the same structure for both chains, but WAT1 is ionized to ^-^OH, and Gln143 is in the canonical amide form (**Fig. 6d**). Residues Tyr34, His30, and Tyr166 are in their neutral forms with a SSHB present between Tyr166 and His30 at a distance of 1.7 Å. Overall, our data indicate that a H_2_O_2_ molecule undergoes deprotonation prior to binding Mn^2+^, and His30, Tyr166, Gln143, and Tyr34 are capable of changing protonation states within the same divalent redox state that coincides with the presence of several SSHBs.

**Fig. 6:**
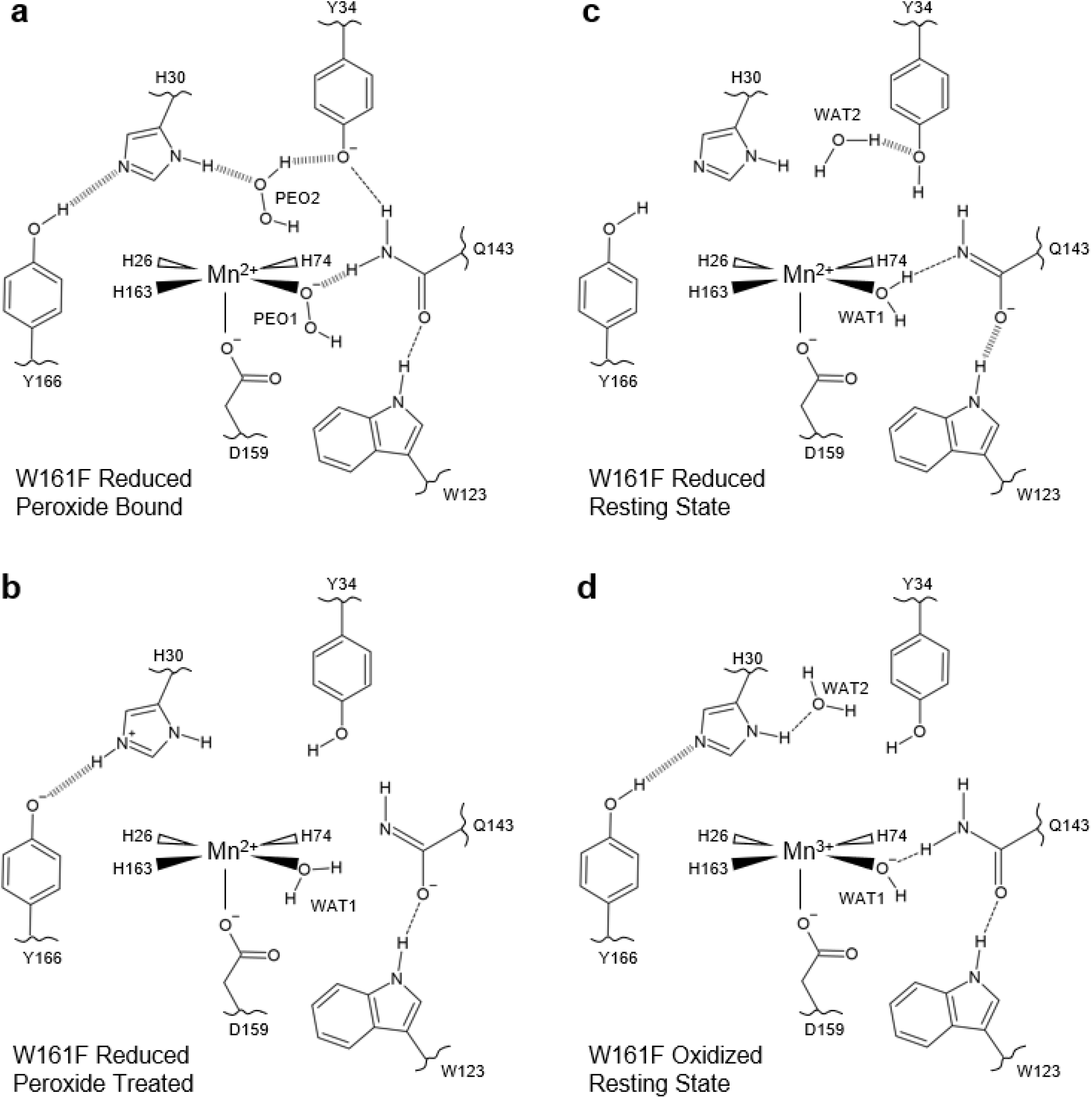
Summary of active site configurations observed for D_2_O_2_-soaked, reduced, and oxidized Trp161Phe MnSOD. a. Dioxygen-bound Trp161Phe MnSOD. The singly-protonated dioxygen species bound to the metal is probably hydroperoxyl anion as supported by DFT calculations. **b** Active site of Trp161Phe MnSOD treated with D_2_O_2_, though cryocooling did not capture a dioxygen species. **c** Reduced resting state of Trp161Phe MnSOD. **d** Oxidized resting state of Trp161Phe MnSOD. Dashed lines represent normal hydrogen bonds and wide dashed lines are SSHBs. The portrayal of structures and bond lengths in 2D are not representative of those seen experimentally in 3D.

Altogether, the present work reveals key features of product inhibition and the related active site proton environment. Through combined neutron diffraction, XAS, and computational chemistry calculations, we find that (1) product inhibition may be initiated from already formed H_2_O_2_ entering the active site; (2) H_2_O_2_ treatment reduces the active site Mn ion; (3) the inhibited complex is a five-coordinate Mn^2+^ complex where a HO_2_^-^ replaces WAT1 found in the resting states; (4) how easily WAT1 may be displaced correlates with the extent of product inhibition; (5) preference for dioxygen species binding in place of the WAT1 position is a result of the overlap with the Mn 3d_z_^2^ orbital; (6) H_2_O_2_ strongly binds to an anionic Tyr34 and His30; (7) an anionic Tyr166 and cationic His30 forms at least transiently with H_2_O_2_ treatment, and (8) slightly different orientations of active site residues can alter their pK_a_s due to the strong electrostatic vectors provided by the Mn metal and limited solvent accessibility. With this new knowledge of product binding, we propose a mechanism for how product inhibition and relief may occur.

### Mechanism for MnSOD product inhibition and relief

Our data are best described by a model where H_2_O_2_ enters the active site in tandem with O ^●-^ donating an electron to Mn^3+^ to form a Mn^2+^-inhibited complex. This is in line with previous observations where inhibition is initiated from the introduction of H_2_O_2_ with Mn^3+^SOD and not Mn^2+^SOD^45,46^. The O_2_^●-^ generation from mixing H_2_O_2_ with MnSOD was explained by Bull *et al.* in 1991 and Hearn *et al.* in 1999, where a back-reaction produces O_2_^●-46,72^. Building on these foundations, our model of product inhibition begins with a H_2_O_2_ molecule entering the active site of Mn^3+^SOD and coordinating between His30 and anionic Tyr34 via SSHBs (**Fig. 7a**). Inhibition is initiated when Mn^3+^ acquires an electron from O_2_^●-^ and a proton is removed from H_2_O_2_ by Tyr34, though His30 has also been shown to change protonation states with either of its nitrogen atoms and could also potentially extract a proton^31^. After the PCET, HO_2_^-^ immediately displaces WAT1 to bind the Mn^2+^ ion and forms a SSHB with Gln143 to make the inhibited complex (**Fig. 7b**). Note that reaction with O ^●-^ is represented by a gain of an electron due to the lack of experimental evidence for O_2_^●-^ binding and uncertainty in whether it requires coordination to the Mn ion for redox catalysis. Decay of the inhibited complex involves a Gln143 to HO_2_^-^ proton transfer that yields a Gln143 in the amide anion form, and the H_2_O_2_ product is subsequently replaced by a water molecule (**Fig. 7c, d**).

**Fig. 7:**
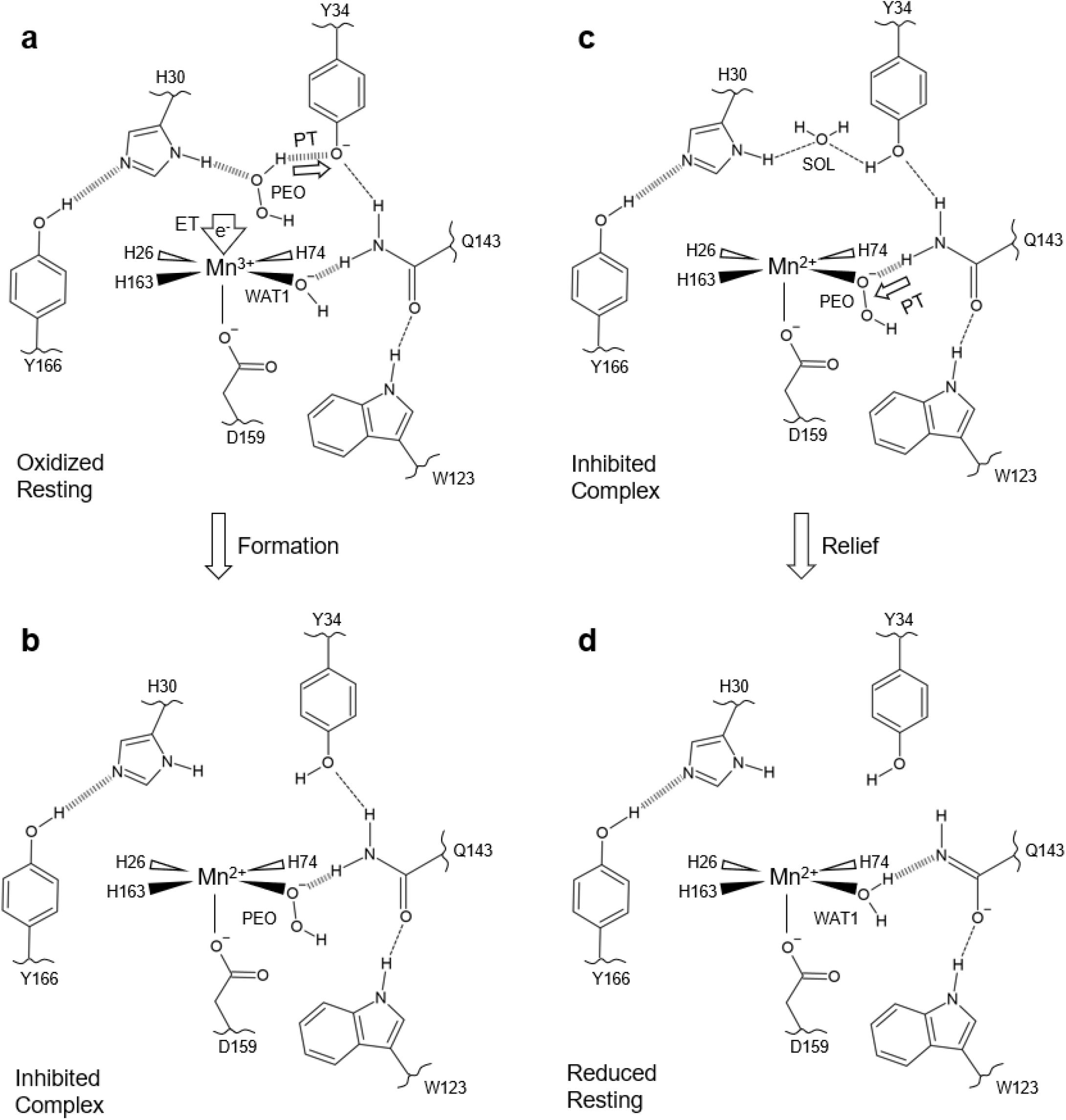
A suggested mechanism for MnSOD product inhibition and relief. a. Product inhibition is dependent on the presence of H_2_O_2_ (denoted as PEO) coordinated between His30 and Tyr34 during the Mn^3+^ → Mn^2+^ redox transition. Due to the lack of experimental evidence for O_2_^●-^ binding and uncertainty in whether it requires coordination with the Mn ion for redox catalysis, the redox reaction is instead represented by a gain of an electron. For the formation of the inhibited complex to proceed, the gain of an electron by Mn^3+^ coincides with the deprotonation of H_2_O_2_ by Tyr34. Note that His30 has been shown to change protonation states on both of its nitrogen atoms and could potentially extract a proton from H_2_O_2_ instead of Tyr34. **b** After the PCET, HO_2_^-^ replaces the WAT1 solvent molecule to form the inhibited complex characterized by the elimination of a Gln143-WAT1 interaction while the Mn ion is in the divalent redox state. **c** The relief of the inhibited complex involves protonation of HO_2_^-^ by Gln143 to form H_2_O_2_ and an ionized Gln143 and subsequent replacement of the original WAT1 position by a water molecule. **d** After H_2_O_2_ leaves the active site, the Mn^2+^SOD is formed that is characterized by an ionized Gln143 forming a SSHB with WAT1, and Tyr34, His30, and Tyr166 in the neutral states. Dashed lines represent normal hydrogen bond and wide dashed lines are SSHBs. The portrayal of the displayed structures and bond lengths in 2D are not representative of those seen experimentally in 3D.

Altogether, the proposed mechanism of product inhibition is dependent on the presence of H_2_O_2_ within the active site during the Mn^3+^ to Mn^2+^ redox transition, an H_2_O_2_ deprotonation event by Tyr34 or His30, and subsequent displacement of WAT1 by HO_2_^-^. Inhibition is characterized by the elimination of the back-and-forth WAT1 and Gln143 proton transfer that is central for MnSOD PCET catalysis^31,59^, and the dioxygen species replacing the WAT1 position is a result of favored molecular orbital overlap with the Mn 3d_z_^2^ α orbital. Relief of the complex involves the protonation of HO_2_^-^ to form H_2_O_2_, which is replaced by a water molecule. The combination of neutron diffraction and XAS has revealed, to our knowledge, the first direct experimental evidence for the identity of the inhibited complex and how it is formed and relieved. Most published mechanistic models of MnSOD product inhibition have presumed that inhibition proceeds through a Mn^2+^SOD and O_2_^●-^ reaction (*k*_3_, **Table 1**)^25–27^, without the involvement of a Gln143 proton transfer, and independent of the Mn^3+^ → Mn^2+^ half-reaction, though our data are best described by the contrary. We also show with cryo-neutron crystallography that Tyr166 and His30 are unambiguously capable of being ionized (**Fig. 5b**), which presents further evidence that the active site metal environment leads to unusual pK_a_s and biochemical states. In total, we conclude that human MnSOD achieves product inhibition through an unusual PCET mechanism. Higher product inhibition in human MnSOD compared to its prokaryotic counterparts likely arose evolutionarily to prevent harmful amounts of H_2_O_2_ from being generated within the mitochondria^26,27^. Abnormalities in mitochondrial H_2_O_2_ concentrations are hallmarks of disease^43,44^, and the precise arrangement of human MnSOD active site residues regulate the output of H_2_O_2_ to prevent mitochondrial dysfunction.

## METHODS

### Perdeuterated expression and purification

For deuterated protein expression of MnSOD, the pCOLADuet-1 expression vector harboring full-length cDNA of *MnSOD* was transformed into *Escherichia coli* BL21(DE3) cells. Transformed cells were grown in D_2_O minimal media within a bioreactor vessel using D_8_-glycerol as the carbon source^73^. Induction was performed with 1 mM isopropyl ꞵ –D-thiogalactopyranoside, 8 mM MnCl_2_, and fed D_8_-glycerol until an OD_600_ of 15.0. Expression was performed at 37 °C for optimal Mn metal incorporation^74^. Harvested cell pastes were stored at –80 °C until purification. For protein purification (with hydrogenated reagents), cells were resuspended in a solution of 5 mM MnCl_2_ and 5 mM 3-(*N*-morpholino)propanesulfonic acid (MOPS), pH 7.8. Clarified lysate was incubated at 55 °C to precipitate contaminant proteins that were subsequently removed by centrifugation. Next, soluble protein was diluted with an equal volume of 50 mM 2-(*N*-morpholino)ethanesulfonic acid (MES) pH 5.5, yielding a final concentration of 25 mM. Measurement of pH verified a value of 5.5 after dilution. Protein was applied onto a carboxymethyl sepharose fast flow column (GE Healthcare) and eluted with a sodium chloride gradient that contained 50 mM MES pH 6.5.

### Crystallization

Perdeuterated MnSOD crystals were grown in either a microgravity environment aboard the International Space Station (ISS) or in an earthly environment at the UNMC Structural Biology Core Facility with hydrogenated reagents. For microgravity crystal growth, crystals were grown in Granada Crystallization Boxes (GCBs, Triana) through capillary counterdiffusion using fused quartz capillary tubes (VitroCom) that had inner diameters of 2.0 mm and outer diameters of 2.4 mm^75^. Crystals were grown from a 25 mg ml^-1^ protein-filled capillary that was plugged with 40 mm of 2% agarose (*w*/*w*) and inserted into a GCB subsequently filled with precipitating agent consisting of 4 M potassium phosphate, pH 7.8. The pH of the phosphate buffer was achieved through 91:9 ratios of K_2_HPO_4_:KH_2_PO_4_. The GCBs were delivered to the ISS by SpX-17 as part of the *Perfect Crystals* NASA payload and returned to earth 1 month later on SpX-18. The crystals within GCBs were observed to be resilient against travel damage and were placed within carry-on baggage during further aircraft travels to the UNMC Structural Biology Core Facility and ORNL. Further details of microgravity crystallization were described previously^76^.

For earthly crystal growth, crystallization was performed using a 9-well glass plate and sandwich box setup (Hampton Research), and the reservoir solution consisted of 1.9 M potassium phosphate adjusted to pH 7.8 by varying ratios of KH_2_PO_4_ and K_2_HPO_4_. The crystallization drop was a mixture of 60 µL of 23 mg mL^-1^ concentrated protein solution (within a buffer of 50 mM MES pH 6.5) and 40 µL of the reservoir solution. Crystals grew up to 0.5 mm^3^ after 6 weeks at 23 °C.

### Crystal Manipulations

Initial deuterium exchange was performed one of two ways, depending on the crystal growth conditions. For microgravity-grown crystals, samples were placed in 1 mL of hydrogenated 4 M potassium phosphate pH 7.8. Deuterium was introduced with 0.1 mL incremental additions every 2 min of 4 M deuterated potassium phosphate (K_2_DPO_4_:KD_2_PO_4_) pD 7.8 (calculated by adding 0.4 to the measured pH reading) for a total of five times and a net volume addition of 0.5 mL. After 10 min, 0.5 mL of the solution was removed leading to a 1 mL solution consisting of 33% deuterium. The process is repeated enough times to gradually increase the deuterium content to ∼100%. The 4 M deuterated potassium phosphate also served as the cryoprotectant for the cryocooling process. Further details of the process were published^51^. For the crystals grown on earth, the initial deuterium exchange of crystals was performed by vapor diffusion in quartz capillaries using deuterated solutions of 2.3 M deuterated potassium phosphate pD 7.8. For cryoprotection, the concentration of the deuterated potassium phosphate was incrementally increased within the capillaries until a concentration of 4 M was achieved.

For redox manipulation, the deuterated potassium phosphate solutions were supplemented with either 6.4 mM potassium permanganate (KMnO_4_) to achieve the Mn^3+^ oxidation state or 300 mM sodium dithionite (Na_2_S_2_O_4_) to achieve the Mn^2+^ state. Crystals were either sealed in capillaries or in 9-well glass plates to ensure maintenance of the desired oxidation state. For the Trp161Phe structure soaked with D_2_O_2_, redox reagents were not used. The dioxygen-bound complex was achieved by supplementing the cryoprotectant that the crystal was immersed in with D_2_O_2_ at a final concentration of 1% (*v*/*v*) and soaking for 5 min before cryocooling. Flash-cooling was performed with an Oxford diffraction cryostream^77^. Further details of ligand cryotrapping were published^51^.

### Crystallographic Data Collection

Time-of-flight, wavelength-resolved neutron Laue diffraction data were collected from perdeuterated crystals using the MaNDi instrument^78,79^ at the Oak Ridge National Laboratory Spallation Neutron Source with wavelengths between 2 to 4 Å. Sample sizes ranged from 0.3 to 0.6 mm^3^ and data were collected to 2.30 Å resolution or better (**Supplementary Table 4**). Crystals were held in stationary positions during diffraction and successive diffraction frames were collected along rotations of the Φ axis. X-ray diffraction data were collected from crystals grown in conditions identical to those used for neutron diffraction using our Rigaku FR-E SuperBright home source (**Supplementary Table 4**).

### Crystallographic Data Processing and Refinement

Neutron data were integrated using the MANTID software package^80–82^ and wavelength-normalized and scaled with LAUENORM from the Daresbury Laue Software Suite^83^. X-ray diffraction data were processed using HKL-3000^84^. Refinements of both neutron and X-ray models were completed separately with PHENIX.REFINE from the PHENIX suite^85^. The refinements were intentionally performed separately due to the known perturbations that X-rays have on the solvent structure, metal redox state, and metal coordination^32,86^. The X-ray model was first refined against its corresponding data set and subsequently used as the starting model for neutron refinement. Torsional backbone angle restraints were derived from the X-ray model and applied to neutron refinement using a geometric target function with PHENIX.REFINE^85^. The neutron refinement process was performed to model the D atoms of the active site last to limit phase bias. Initial rounds of refinement to fit protein structure included only non-exchangeable D atoms which have stereochemical predictable positions. Afterward, H/D atoms were modeled onto the position of each amide proton, and occupancy was refined. In general, the asymmetric units of the neutron crystal structures had a deuterium content of ∼85% for the amide backbone, and areas with low deuterium exchange (< 50 %) coincided with the presence of hydrogen bonds forming a secondary structure. Next, exchangeable proton positions of residues outside the active site (e.g. hydroxyl group of serine/tyrosine) were manually inspected for obvious positive omit |*F*_o_| – |*F*_c_| neutron scattering length density at a contour of 2.5σ or greater and modeled as a fully occupied deuterium. If the density was not obvious, and there was no chemically sensible reason for the residue to be deprotonated (which is the case for residues outside the active site), the proton position was H/D occupancy refined. D_2_O molecules outside the active site were then modeled and adjusted according to the nuclear density. Last, D atoms of the active site were modeled manually. At the active site, a residue is considered deprotonated when (1) attempts to model and refine a proton result in negative |*F*_o_| – |*F*_c_| difference neutron scattering length density, (2) all the other protons of the residue can be placed, and (3) the heavy atom that is deprotonated acts as a hydrogen-bond acceptor.

### X-ray Absorption Spectroscopy Measurements

Mn K-edge HERFD-XANES spectra were recorded at beamline 15-2 of the Stanford Synchrotron Radiation Lightsource (SSRL) while Mn K-edge EXAFS spectra were collected at beamline 9-3. At both beamlines, data were collected at 10 K using a liquid He cryostat, and the incident energy was tuned to the first derivative of an internal Mn foil at 6539 eV. X-ray irradiation was carefully monitored so that two subsequent scans of the same spot did not have photoreduction differences and different spots along samples were scanned. When appropriate, aluminum foil was inserted into the beam path to attenuate the incident flux. For HERFD-XANES measurements, a Johann-type hard X-ray spectrometer with six Ge(333) analyzer crystals was used with a liquid-nitrogen cooled Si(311) double crystal monochromator, and energy was calibrated to a glitch with measurement of Mn Foil. For EXAFS, measurements were recorded with a 100-element Ge monolithic solid-state fluorescence detector and a Si(220) monochromator at Φ = 90° was used.

### X-ray Absorption Spectroscopy Data Analysis

Pre-edge peak fitting for HERFD results was performed with pseudo-Voigt functions where the post-edge of the XANES is normalized to unity. Pre-edge intensities are defined as the total trapezoidal numerical integration of the fitted peaks. EXAFS data reduction, averaging, and refinement were carried out with the LARCH software package^87^. Refinement of the *k*^2^χ(*k*) EXAFS data used phases and amplitudes obtained from FEFF^88^. For each fit, optimization of the radial distribution around the absorbing Mn ion (*r*) and the Debye-Waller factor (σ^2^) was performed. The goodness-of-it was evaluated by reduced χ^2^ values and R-factors.

### Computational Methods

All DFT calculations were performed with the ORCA quantum chemistry package version 5.0 using the B3LYP functional, the def2-TZVP basis set for all atoms, and the CPCM solvation model^89–92^. For geometry optimizations, the full active site (i.e., residues shown in the right panel of **Fig. 1a**) was included from the neutron structure where the O and N atoms of the peptide backbone were truncated and C^α^ was fixed. Additional fixed restraints were placed on aromatic residues found on the periphery of the active site (Phe66, Trp123, Trp161, and Tyr166) to mimic the packing found in the native enzyme. The Mn ion used the high-spin quintet and sextet states for trivalent and divalent systems, respectively, per experimental observations^66^. A dense integration grid and tight convergence were enforced.

For TD-DFT calculations, the Mn ion was instead assigned the core property basis set, CP(PPP)^93,94^. The geometry optimized model was used and truncated to only the Mn ion and its immediate ligands. Inclusion of all active site residues for TD-DFT did not significantly alter the simulated spectra. Computed Mn K pre-edge data were plotted using a Gaussian broadening of 1 eV and a 32.6 eV energy correction was applied in line with previous studies^61,68^.

XANES simulations of the Mn K-edge were achieved using the FDMNES software package^95^. The scattering potential around the Mn absorber was calculated self-consistently within a radius of 6 Å and a fully screened core-hole using the finite difference method was used. XANES spectra were simulated for Mn complexes derived from the neutron structural data and fit to experimental HERFD-XANES spectra using the Fitit code^65^. For fits of the spectra, the XANES spectra of the neutron structure is first simulated with the finite difference method (a DFT approach)^95^. Systematic deformations of the structure were then applied, and the corresponding spectra of each deformation were simulated to generate a training set. From the training set, each point of a XANES spectrum is defined as a function of structural parameters that are then used for fitting the experimental spectra with structural refinement. Fits were performed by refinement of the Mn bond distances.

## DATA AVAILABILITY

Coordinates and structure factors for neutron and X-ray crystallographic data presented in this study have been deposited in the Protein Data Bank (PDB 8VHW [https://doi.org/10.2210/pdb8vhw/pdb], PDB 8VHY [https://doi.org/10.2210/pdb8vhy/pdb], PDB 8VJ0 [https://doi.org/10.2210/pdb8vj0/pdb], PDB 8VJ4 [https://doi.org/10.2210/pdb8vj4/pdb], PDB 8VJ5 [https://doi.org/10.2210/pdb8vj5/pdb], and PDB 8VJ8 [https://doi.org/10.2210/pdb8vj8/pdb]). All relevant data supporting the key findings of this study are available within the article and its Supplementary Information files or from the corresponding author upon reasonable request.

## ACKNOWLEDGEMENTS

This research was supported by the NIH (R01-GM145647) and NASA EPSCoR (NE-80NSSC17M0030 and NE-NNX15AM82A). The UNMC Structural Biology Core Facility was funded by the Fred and Pamela Buffett NCI Cancer Center Support Grant (P30CA036727). The research at Oak Ridge National Laboratory (ORNL) Spallation Neutron Source was sponsored by the Scientific User Facilities Division, Office of Basic Energy Sciences, US Department of Energy. The Office of Biological and Environmental Research supported research at ORNL Center for Structural Molecular Biology (CSMB) using facilities supported by the Scientific User Facilities Division, Office of Basic Energy Sciences, US Department of Energy. Use of the Stanford Synchrotron Radiation Lightsource (SSRL), SLAC National Accelerator Laboratory, is supported by the US Department of Energy (DOE), Office of Science, Office of Basic Energy Sciences under Contract DE-AC02-76SF00515. The SSRL Structural Molecular Biology Program is supported by the DOE Office of Biological and Environmental Research, and by the National Institutes of Health, National Institute of General Medical Sciences (P30GM133894). The contents of this publication are solely the responsibility of the authors and do not necessarily represent the official views of NIGMS or NIH. Quantum chemical computations were completed using the Holland Computing Center of the University of Nebraska, which receives support from the Nebraska Research Initiative.

