## Supplemental Information for "Revealing the atomic and electronic mechanism of human manganese superoxide dismutase product inhibition"

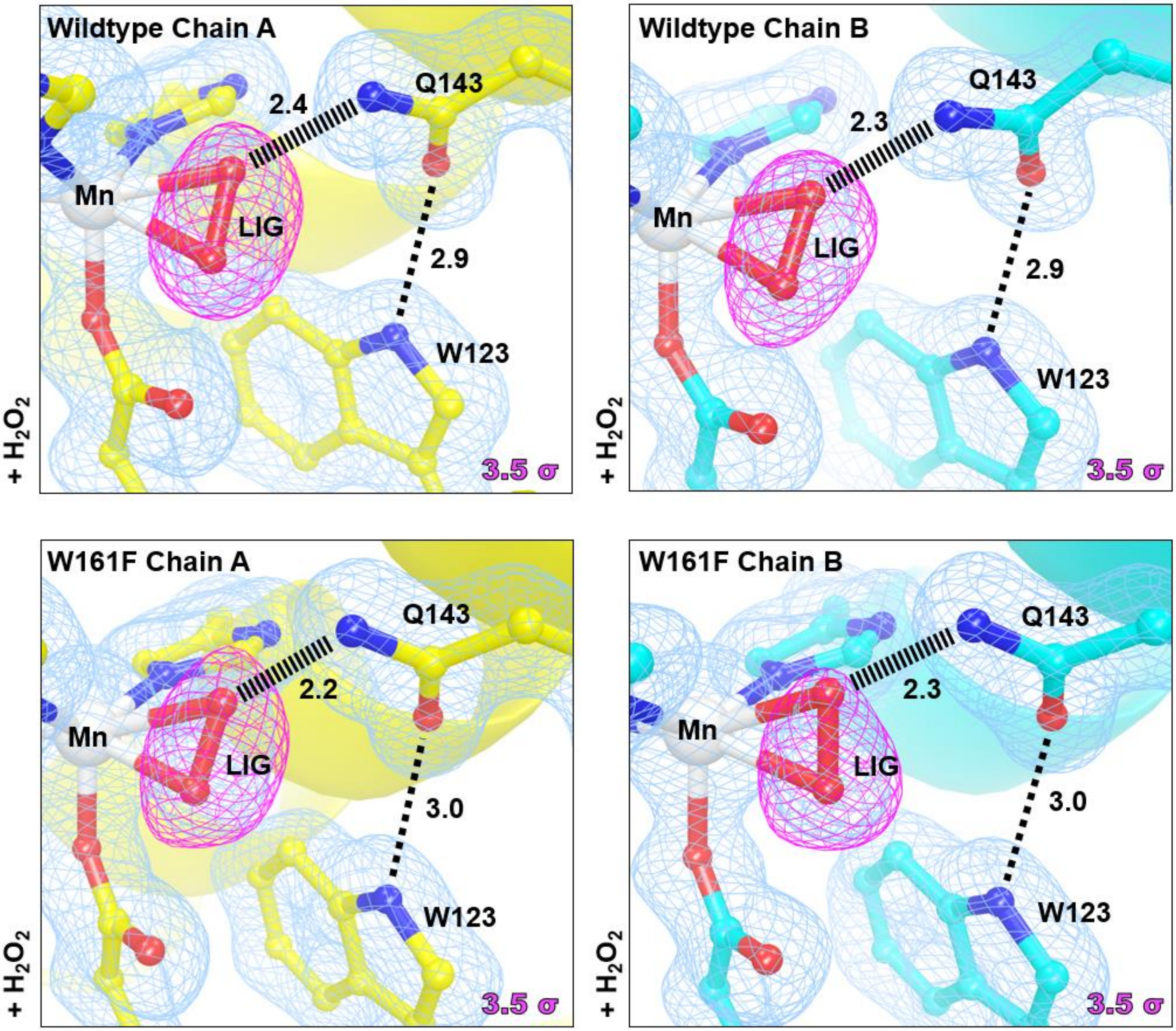

**Supplementary Figure 1. X-ray structures of Wildtype and Trp161Phe MnSOD soaked with $H_2O_2$.** Magenta omit $|F_o| - |F_c|$ difference electron density is displayed at 3.5σ. Light blue $2|F_o| - |F_c|$ density is displayed at 1.0σ. Distances are in Å. Dashed lines indicate typical hydrogen bonds and hashed lines indicate SSHBs that are hydrogen bonds < 2.8 Å. Note that due to photoreduction effects, the dioxygen species are refined at partial occupancy and have side-on binding orientations compared to the neutron structure counterpart that is absent of photoreduction and was refined at full occupancy and has an end-on binding orientation.

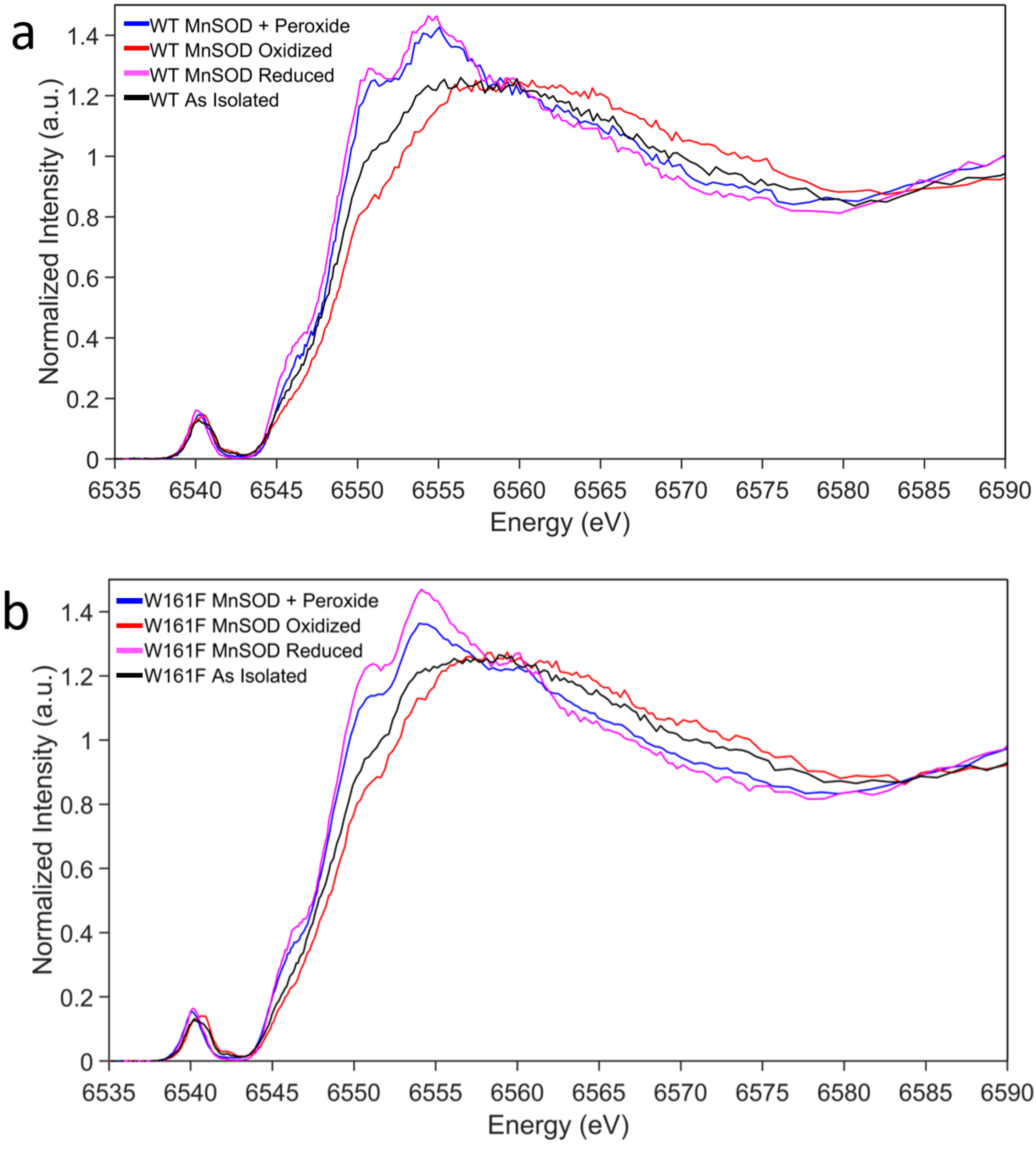

**Supplementary Figure 2. Kα HERFD-XANES spectra of MnSOD**. **a** HERFD-XANES of wildtype MnSOD in the peroxide-soaked, oxidized, reduced, and as isolated forms. **b** HERFD-XANES of Trp161Phe MnSOD in the peroxide-soaked, oxidized, reduced, and as isolated forms. The oxidized and reduced samples correspond to $Mn^{3+}$SOD and $Mn^{2+}$SOD resting states.

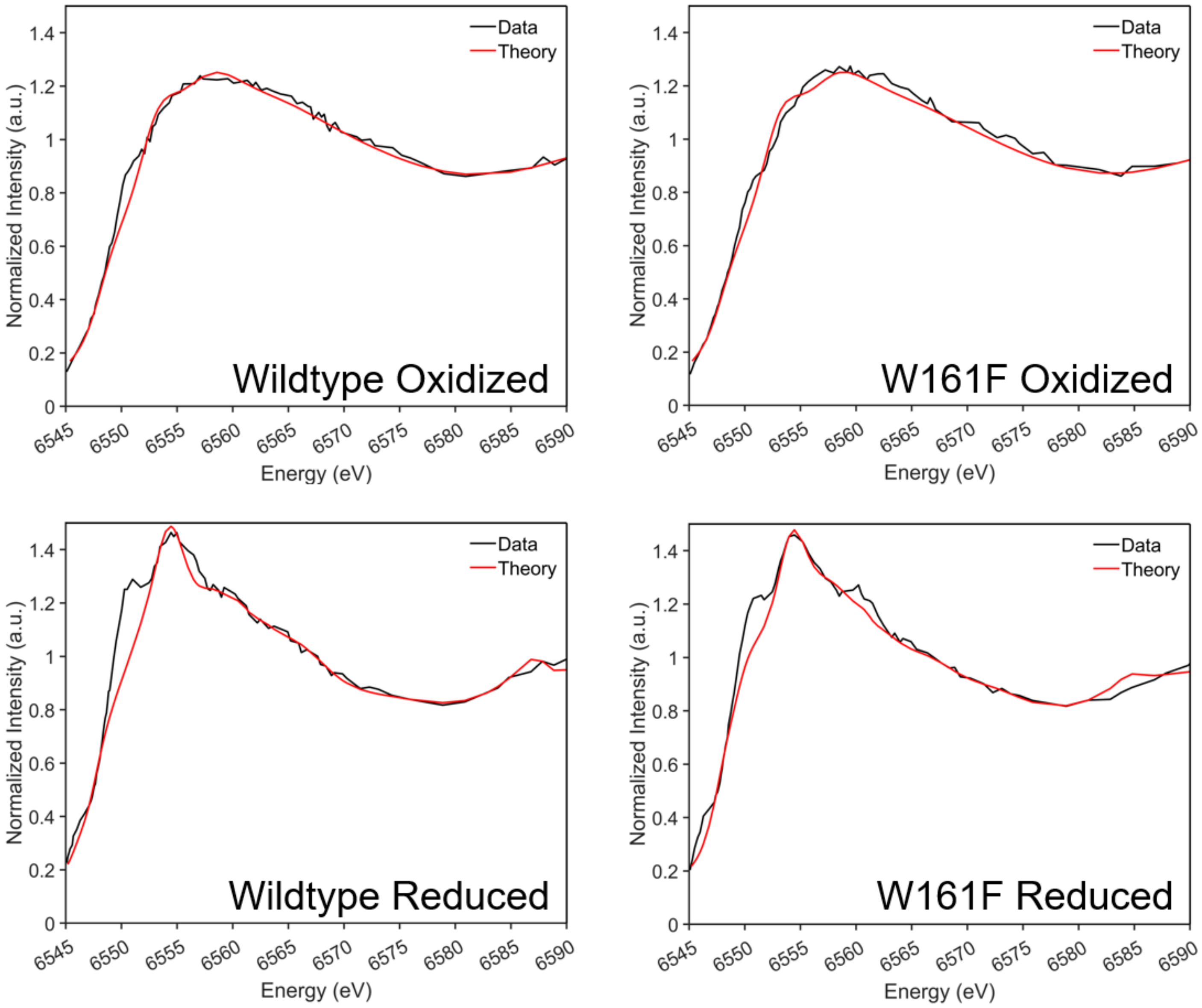

**Supplementary Figure 3. Fit of oxidized and reduced HERFD-XANES spectra for wildtype and Trp161Phe MnSOD**. Oxidized and reduced samples correspond to $Mn^{3+}$SOD and $Mn^{2+}$SOD resting states. Spectra were simulated with FDMNES, and fits were performed using FITIT code.

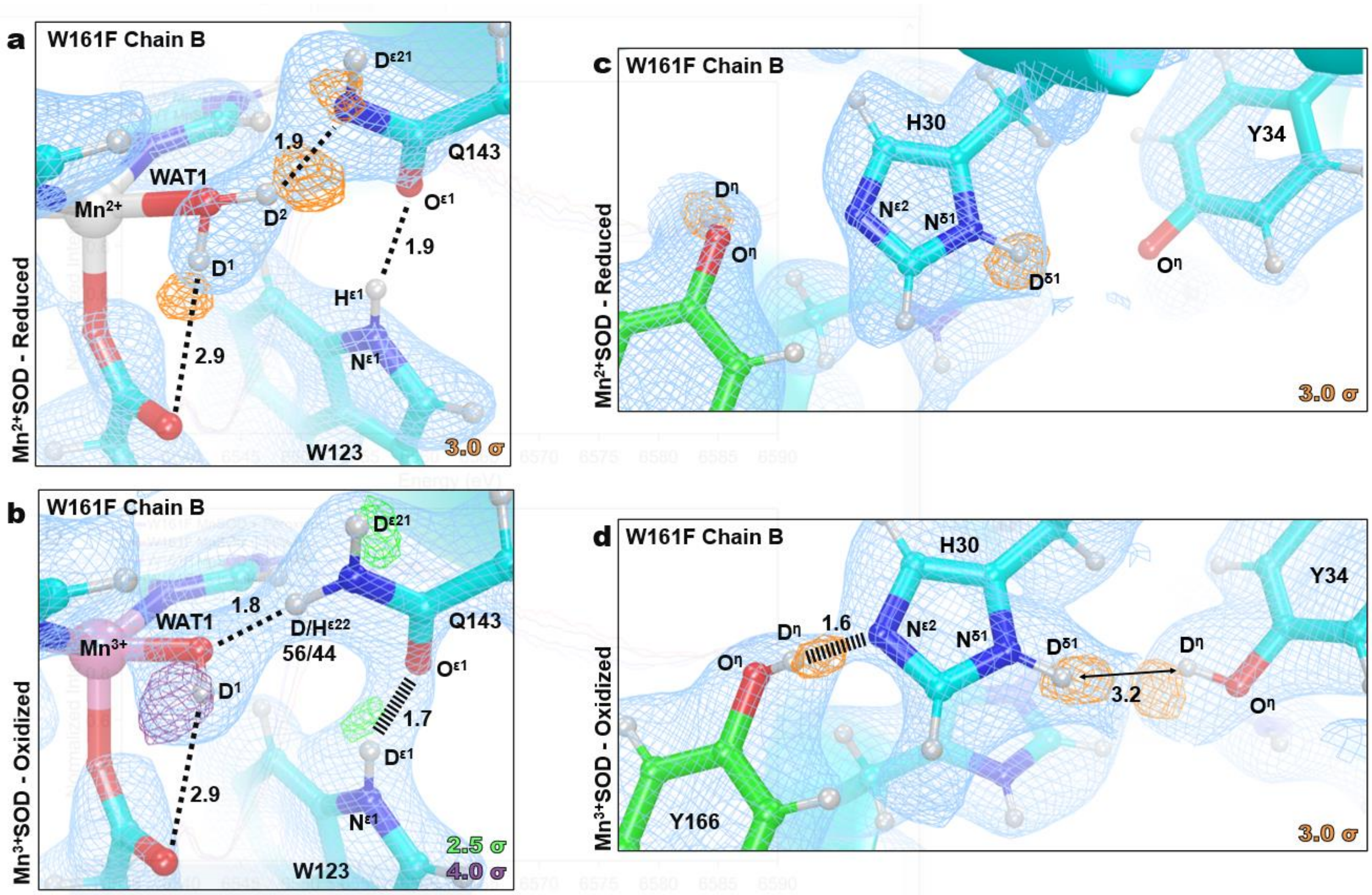

**Supplementary Figure 4. Neutron Structure and protonation states at the active site of Trp161Phe $Mn^{2+}$SOD and Trp161Phe $Mn^{3+}$SOD.** **a** Neutron structure of Trp161Phe $Mn^{2+}$SOD at the active site of chain B with orange omit $|F_o|$ - $|F_c|$ difference neutron scattering length density of protons displayed at 3.0σ. **b** Neutron structure of Trp161Phe $Mn^{3+}$SOD at the active site of chain B with green and purple omit $|F_o|$ - $|F_c|$ difference neutron scattering length density of protons displayed at 2.5σ and 4.0σ, respectively. For the $D^{\varepsilon 22}$(Q143), positive omit difference density was not present above 2.0σ due to density cancellation with the negative neutron scattering length density of hydrogen. As an alternative, the proton position was occupancy refined to yield a ratio of 56% deuterium and 44% hydrogen. **c** Neutron structure of second-sphere residues at the active site of Trp161Phe $Mn^{2+}$SOD chain B with orange omit $|F_o|$ - $|F_c|$ difference neutron scattering length density of protons displayed at 3.0σ. Due to a lack of interpretable density for Tyr34, a protonation state was not definitively assigned. **d** Neutron structure of second-sphere residues at the active site of Trp161Phe $Mn^{3+}$SOD chain B with orange omit $|F_o|$ - $|F_c|$ difference neutron scattering length density of protons displayed at 3.0σ. Light blue $2|F_o|$ - $|F_c|$ density is displayed at 1.0σ. Distances are in Å. Dashed lines indicate typical hydrogen bonds, and hashed lines indicate SSHBs, hydrogen bonds < 1.8 Å.

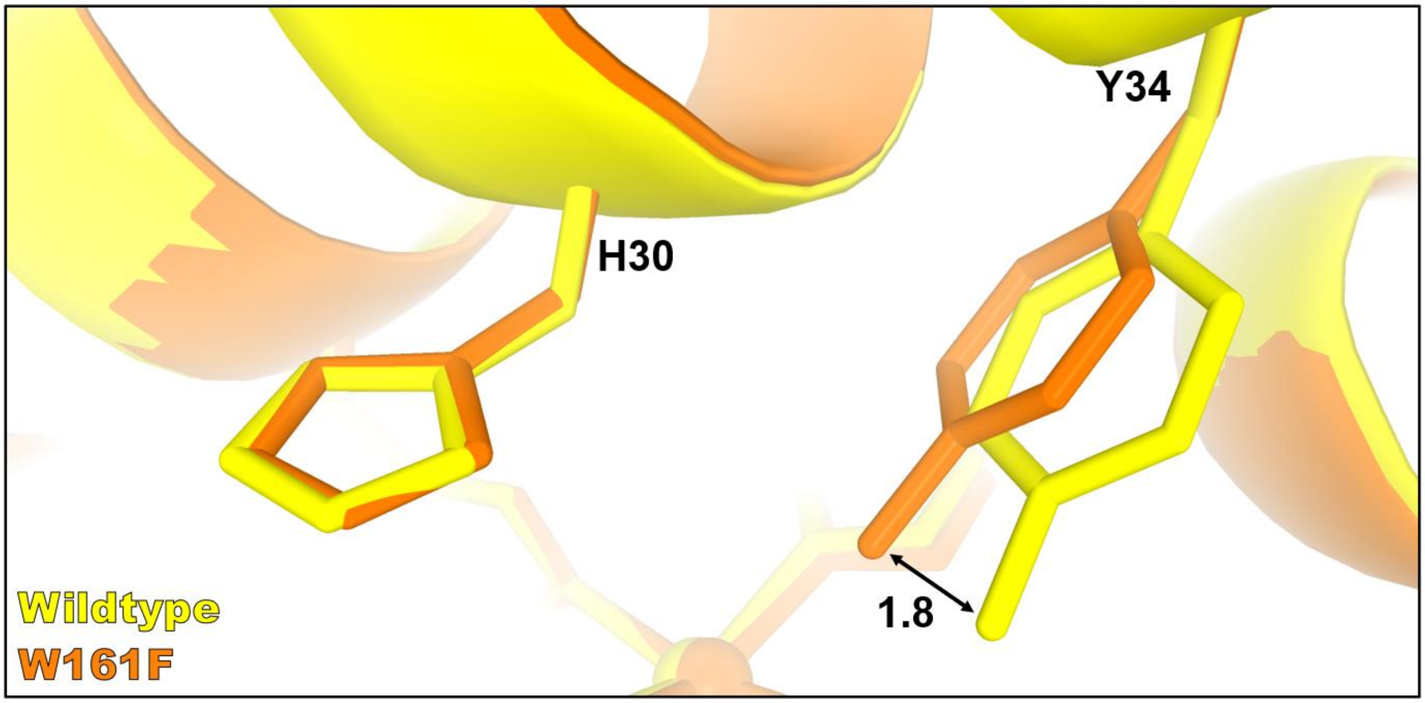

**Supplementary Figure 5. Movement of Tyr34 in the Trp161Phe MnSOD variant.** Active site overlay of wildtype (yellow) and Trp161Phe (orange) MnSOD highlighting a 1.8 Å movement of the Tyr34 hydroxyl group.

**Supplementary Table 1. Active Site Mn Bond Lengths of MnSOD Neutron Structures.**

| | Trp161Phe $D_2O_2$-Soaked[b] | | Trp161Phe $Mn^{3+}$SOD | | Trp161Phe $Mn^{2+}$SOD | | Wildtype $Mn^{3+}$SOD | | Wildtype $Mn^{2+}$SOD[c] | |
|---|---|---|---|---|---|---|---|---|---|---|
| Mn Bonds (Å) | A | B | A | B | A | B | A | B | A | B |
| Mn-$N^{\varepsilon 2}$(H26) | 2.13 | 2.20 | 2.01 | 2.01 | 2.20 | 2.11 | 2.07 | 2.07 | 2.26 | 2.10 |
| Mn-$N^{\varepsilon 2}$(H74) | 2.13 | 2.28 | 2.15 | 2.15 | 2.19 | 2.16 | 2.13 | 2.12 | 2.19 | 2.25 |
| Mn-$O^{\varepsilon 2}$(D159) | 2.20 | 2.05 | 2.01 | 2.01 | 2.13 | 2.20 | 1.95 | 1.94 | 2.44 | 2.15 |
| Mn-$N^{\varepsilon 2}$(H163) | 2.25 | 2.17 | 2.15 | 2.15 | 2.14 | 2.15 | 2.06 | 2.14 | 2.23 | 2.21 |
| Mn-O(WAT1) | 2.13 | - | 1.84 | 1.84 | 2.37 | 2.24 | 1.78 | 1.76 | 2.12 | 2.22 |
| Mn-$O^1$(LIG)[a] | - | 1.94 | - | - | - | - | - | - | - | - |
| Mn-O(OL) | - | - | - | - | - | - | - | - | 1.82 | - |

[a]$O^1$(LIG) refers to the closest oxygen atom of the dioxygen species.

[b]Only chain B of $D_2O_2$-soaked Trp161Phe MnSOD is bound by a dioxygen species, denoted as LIG. Chain A is in the typical five-coordinated state bound by WAT1.

[c]For chain A of wildtype $Mn^{2+}$SOD, an $^-$OD molecule is observed binding opposite of Asp159 and is six-coordinate. Chain B is in the typical five-coordinated state.

**Supplementary Table 2. EXAFS Fitting Results for $H_2O_2$-soaked Trp161Phe MnSOD.**

| Mn-O | | | Mn-N | | | Mn···C | | | Mn···O | | | Mn···O···O | | |
|---|---|---|---|---|---|---|---|---|---|---|---|---|---|---|
| *n* | *r* (Å) | $\sigma^2$ x $10^3$ (Å$^2$) | *n* | *r* (Å) | $\sigma^2$ x $10^3$ (Å$^2$) | *n* | *r* (Å) | $\sigma^2$ x $10^3$ (Å$^2$) | *n* | *r* (Å) | $\sigma^2$ x $10^3$ (Å$^2$) | *n* | *r* (Å) | $\sigma^2$ x $10^3$ (Å$^2$) |
| 2 | 2.04 | 2.5 | 3 | 2.20 | 2.5 | 7 | 3.15 | 10 | 1 | 2.52 | 0 | 2 | 2.63 | 0 |
| Mn···C···O | | | Mn···C···N | | | $\chi^2$ | | | Reduced $\chi^2$ | | | R-Factor | | |
| *n* | *r* (Å) | $\sigma^2$ x $10^3$ (Å$^2$) | *n* | *r* (Å) | $\sigma^2$ x $10^3$ (Å$^2$) | 14.28 | | | 3.56 | | | 0.0197 | | |
| 2 | 3.26 | 0 | 12 | 3.48 | 2.5 | | | | | | | | | |

**Supplementary Table 3. Comparison of MnSOD bond lengths from various methods.**

| **Trp161Phe $Mn^{3+}$SOD** | | | |
|---|---|---|---|
| Bond | Neutron Structure (Å) | DFT (Å) | XANES Fit (Å) |
| Mn-N$^{\varepsilon 2}$(H26) | 2.01 | 2.02 | 1.96 |
| Mn-N$^{\varepsilon 2}$(H74) | 2.15 | 2.11 | 2.04 |
| Mn-N$^{\varepsilon 2}$(H163) | 2.15 | 2.08 | 2.06 |
| Mn-O$^{\delta 2}$(D159) | 2.01 | 1.97 | 1.96 |
| Mn-O(WAT1) | 1.84 | 1.82 | 1.80 |
| **Trp161Phe $Mn^{2+}$SOD** | | | |
| Bond | Neutron Structure (Å) | DFT (Å) | XANES Fit (Å) |
| Mn-N$^{\varepsilon 2}$(H26) | 2.20 | 2.20 | 2.14 |
| Mn-N$^{\varepsilon 2}$(H74) | 2.19 | 2.20 | 2.26 |
| Mn-N$^{\varepsilon 2}$(H163) | 2.14 | 2.19 | 2.11 |
| Mn-O$^{\delta 2}$(D159) | 2.13 | 2.07 | 2.08 |
| Mn-O(WAT1) | 2.37 | 2.11 | 2.37 |
| **Wildtype $Mn^{3+}$SOD** | | | |
| Bond | Neutron Structure (Å) | DFT (Å) | XANES Fit (Å) |
| Mn-N$^{\varepsilon 2}$(H26) | 2.07 | 2.02 | 1.99 |
| Mn-N$^{\varepsilon 2}$(H74) | 2.12 | 2.11 | 2.08 |
| Mn-N$^{\varepsilon 2}$(H163) | 2.14 | 2.09 | 2.09 |
| Mn-O$^{\delta 2}$(D159) | 1.94 | 1.96 | 1.95 |
| Mn-O(WAT1) | 1.76 | 1.82 | 1.80 |
| **Wildtype $Mn^{2+}$SOD** | | | |
| Bond | Neutron Structure (Å) | DFT (Å) | XANES Fit (Å) |
| Mn-N$^{\varepsilon 2}$(H26) | 2.10 | 2.18 | 2.06 |
| Mn-N$^{\varepsilon 2}$(H74) | 2.25 | 2.24 | 2.34 |
| Mn-N$^{\varepsilon 2}$(H163) | 2.21 | 2.22 | 2.25 |
| Mn-O$^{\delta 2}$(D159) | 2.15 | 2.10 | 2.02 |
| Mn-O(WAT1) | 2.22 | 2.12 | 2.25 |

**Supplementary Table 4. Data collection and refinement statistics for MnSOD.**

| Data Collection Statistics | | | | | | |
|---|---|---|---|---|---|---|
| | Neutron | | | X-ray | | |
| Variant | Trp161Phe | Trp161Phe | Trp161Phe | Trp161Phe | Wildtype | Trp161Phe |
| Chemical State | $D_2O_2$-Soaked | Reduced | Oxidized | $H_2O_2$-Soaked | $H_2O_2$-Soaked | Reduced |
| PDB Code | 8VHW | 8VHY | 8VJ0 | 8VJ4 | 8VJ5 | 8VJ8 |
| Diffraction Source | MaNDi | | | Rigaku FR-E SuperBright | | |
| Temperature (K) | 100 | 100 | 296 | 100 | 100 | 100 |
| Space group | $P6_122$ | | | $P6_122$ | | |
| *a*, *b*, *c* (Å) | 77.80, 77.80, 236.80 | 78.11, 78.11, 236.32 | 80.75, 80.75, 239.43 | 77.66, 77.66, 234.20 | 78.35, 78.35, 236.63 | 77.91, 77.91, 235.14 |
| $\alpha, \beta, \gamma$ (°) | 90, 90, 120 | | | 90, 90, 120 | | |
| Wavelengths (Å) | 2-4 | | | 1.5418 | | |
| No. of images | 12 | 11 | 13 | 252 | 202 | 252 |
| Exposure time | 48 h | 36 h | 20 h | 300 s | 300 s | 300 s |
| No. of unique reflections | 19467 | 18253 | 20479 | 48362 | 38708 | 46382 |
| Total No. of reflections | 189145 | 76061 | 122164 | 256135 | 235186 | 456697 |
| Resolution range (Å) | 14.82-2.30 (2.38-2.30) | 14.74-2.30 (2.42-2.30) | 14.89-2.30 (2.42-2.30) | 50.00 – 1.68 (1.72-1.68) | 50.00 – 1.76 (1.80-1.76) | 50.00 – 1.70 (1.74-1.70) |
| Multiplicity | 9.7 (7.0) | 4.2 (3.8) | 6.0 (5.5) | 5.3 (4.1) | 6.1 (3.9) | 9.8 (5.4) |
| I/σ(I) | 9.7 (5.6) | 4.8 (3.1) | 5.9 (3.8) | 11.41 (2.02) | 5.96 (2.24) | 16.51 (1.99) |
| $R_{merge}$ | 0.279 (0.286) | 0.247 (0.306) | 0.202 (0.279) | - | - | - |
| $R_{meas}$ | 0.294 (0.306) | 0.278 (0.347) | 0.220 (0.307) | 0.135 (0.391) | 0.289 (0.602) | 0.122 (0.579) |
| $CC_{1/2}$ | 0.919 (0.607) | 0.840 (0.407) | 0.978 (0.368) | 0.920 (0.769) | 0.872 (0.732) | 0.950 (0.814) |
| $R_{pim}$ | 0.086 (0.102) | 0.122 (0.155) | 0.083 (0.125) | 0.056 (0.194) | 0.104 (0.278) | 0.036 (0.244) |
| Data completeness (%) | 99.0 (98.0) | 92.6 (87.2) | 96.6 (96.4) | 99.3 (97.2) | 88.4 (78.8) | 97.6 (92.5) |
| Refinement Statistics | | | | | | |
| $R_{work}$ | 0.2488 | 0.2752 | 0.2806 | 0.1952 | 0.1993 | 0.1913 |
| $R_{free}$ | 0.2774 | 0.3005 | 0.3089 | 0.2422 | 0.2183 | 0.2237 |
| [a]No. of atoms<br>Protein<br>[b]Solvent<br>Mn | <br>6385<br>725<br>2 | <br>6540<br>1392<br>2 | <br>6228<br>479<br>2 | <br>3168<br>460<br>2 | <br>3174<br>614<br>2 | <br>3168<br>570<br>2 |
| R.m.s. deviations<br>Bond lengths (Å)<br>Bond angles (°) | <br>0.003<br>0.58 | <br>0.002<br>0.53 | <br>0.003<br>0.53 | <br>0.011<br>1.05 | <br>0.003<br>0.58 | <br>0.004<br>0.69 |
| Average *B*-factor<br>Protein<br>Water<br>Mn<br>Peroxide | 20.71<br>21.13<br>17.40<br>19.40<br>21.58 | 11.67<br>11.44<br>13.25<br>7.12<br>- | 17.42<br>17.62<br>13.76<br>9.37<br>- | 19.34<br>18.59<br>24.55<br>14.11<br>13.22 | 21.90<br>20.15<br>30.85<br>14.47<br>14.63 | 16.48<br>15.20<br>23.26<br>10.93<br>- |

[a]No. of atoms is inclusive of H/D atoms for neutron structures.

[b]The neutron structure of oxidized Trp161Phe was collected at room temperature and, as a result, has significantly less observed solvent atoms compared to the other structures.
